# IL-32 is a metabolic regulator promoting survival and proliferation of malignant plasma cells

**DOI:** 10.1101/2021.02.22.431638

**Authors:** Kristin Roseth Aass, Robin Mjelle, Martin H. Kastnes, Synne S. Tryggestad, Luca M. van den Brink, Marita Westhrin, Muhammad Zahoor, Siv H. Moen, Glenn Buene, Kristine Misund, Anne-Marit Sponaas, Qianli Ma, Anders Sundan, Richard WJ Groen, Tobias S. Slørdahl, Anders Waage, Therese Standal

## Abstract

IL-32 is a non-classical cytokine expressed in cancers, inflammatory diseases and infections. IL-32 can have both extracellular and intracellular functions, and its receptor is not identified. We here demonstrate that endogenously expressed, intracellular IL-32 binds to components of the mitochondrial respiratory chain and promotes oxidative phosphorylation. Knocking out IL-32 in malignant plasma cells significantly reduced survival and proliferation *in vitro* and *in vivo*. High throughput transcriptomic and MS-metabolomic profiling of IL-32 KO cells revealed that loss of IL-32 leads to profound perturbations in metabolic pathways, with accumulation of lipids, pyruvate precursors and citrate, indicative of reduced mitochondrial function. IL-32 is expressed in a subgroup of multiple myeloma patients with an inferior prognosis. Primary myeloma cells expressing IL-32 were characterized by a plasma cell gene signature associated with immune activation, proliferation and oxidative phosphorylation. We propose a novel concept for regulation of metabolism by an intracellular cytokine and identify IL-32 as an endogenous growth and survival factor for malignant plasma cells. IL-32 is a potential prognostic biomarker and a treatment target in multiple myeloma.

## Introduction

Multiple myeloma (MM) is a cancer of terminally differentiated plasma cells in the bone marrow^1^. Similar to normal plasma cells, the malignant cells are dependent on the bone marrow microenvironment for survival. Most MM cell growth factors are produced by cells of stromal and hematopoietic origin, and IL-6, APRIL and BAFF are key survival factors. Only a small number of pro-survival or proliferative factors are produced by the cancer cells themselves^1^.

The bone marrow is characterized by areas of low oxygen levels and the master regulator of hypoxic metabolism, HIF1α, is highly expressed in MM cells in hypoxic niches^2–4^. Hypoxic MM cells may exhibit a glycolytic phenotype^4, 5^ but several studies have demonstrated that aerobic metabolism, and thus oxidative phosphorylation (OXPHOS), is fully functional in MM cells. The OXPHOS/glycolysis ratio is dynamic and possibly regulated by microenvironmental cues and state of dormancy.^6–8^ Furthermore, high level of aerobic metabolism may contribute to drug resistance and disease progression in MM^9, 10^.

IL-32 is a pluripotent pro-inflammatory cytokine involved in a range of diseases including cancer, infections and autoimmunity^11, 12^. IL-32 has no sequence homology with other cytokine families, and an IL-32 receptor has not been identified. The secreted form of IL-32 acts like a cytokine, but IL-32 also have distinct intracellular functions^13^. For example, IL-32 was detected in the mitochondrial fraction of breast cancer cells and improved the metabolic fitness of the cells during hypoxic stress^14^. IL-32 is also intriguingly regulated by two different oxygen sensing systems; HIF1α^14, 15^ and cysteamine (2-aminoethanethiol) dioxygenase (ADO)^16^, indicating that this protein has an important function in response to low oxygen tension. We have previously shown that IL-32 is highly expressed in a subgroup of MM patients characterized by poor prognosis and a hypoxic signature and that expression of IL-32 in MM cells is increased in response to hypoxia in a HIF1α-dependent manner.^15^ The role and molecular mechanism of action of IL-32 in plasma cells is however not known.

In this study we demonstrate that endogenous intracellular IL-32 promotes survival and proliferation of MM cells *in vitro* and *in vivo*. IL-32 interacts with components of the mitochondrial respiratory chain and acts as a key regulator of MM cell metabolism. Moreover, IL-32 expression in patient samples is associated with an immature, proliferative plasma cell profile. Our data demonstrate a novel function of IL-32 and support that IL-32 is a potential prognostic biomarker and a treatment target in MM.

## Results

### IL-32 is important for myeloma cell proliferation *in vitro* and tumor engraftment *in vivo*

We have previously demonstrated that IL-32 is expressed by a subgroup of MM cells^15^. Moreover, bone marrow plasma cells obtained from healthy donors express IL-32 at higher levels relative to other B cell subsets (Supplementary Figure 1A). The function of IL-32 in plasma cells is however unknown. To investigate the role of IL-32 in MM cells we depleted IL-32 using CRISPR/Cas9 from three IL-32-expressing cell lines JJN3, INA-6 and H929 (Supplementary Figure 1B). These cell lines have different IgH translocations t(14;16), t(11;14) and t(4;14), respectively, and also differ in terms of p53 and RAS mutations ^17, 18^. INA-6 is one of a few IL-6-dependent MM cell lines and thus more similar to primary MM cells. Strikingly, for all three cell lines, loss of IL-32 significantly reduced proliferation as assessed by automated cell counting (Figure 1A - C). The viability of the INA-6 KO cells was significantly reduced compared with mock (WT) while there was no difference in viability for H929 and JJN3 (Figure 1D).

**Figure 1:**
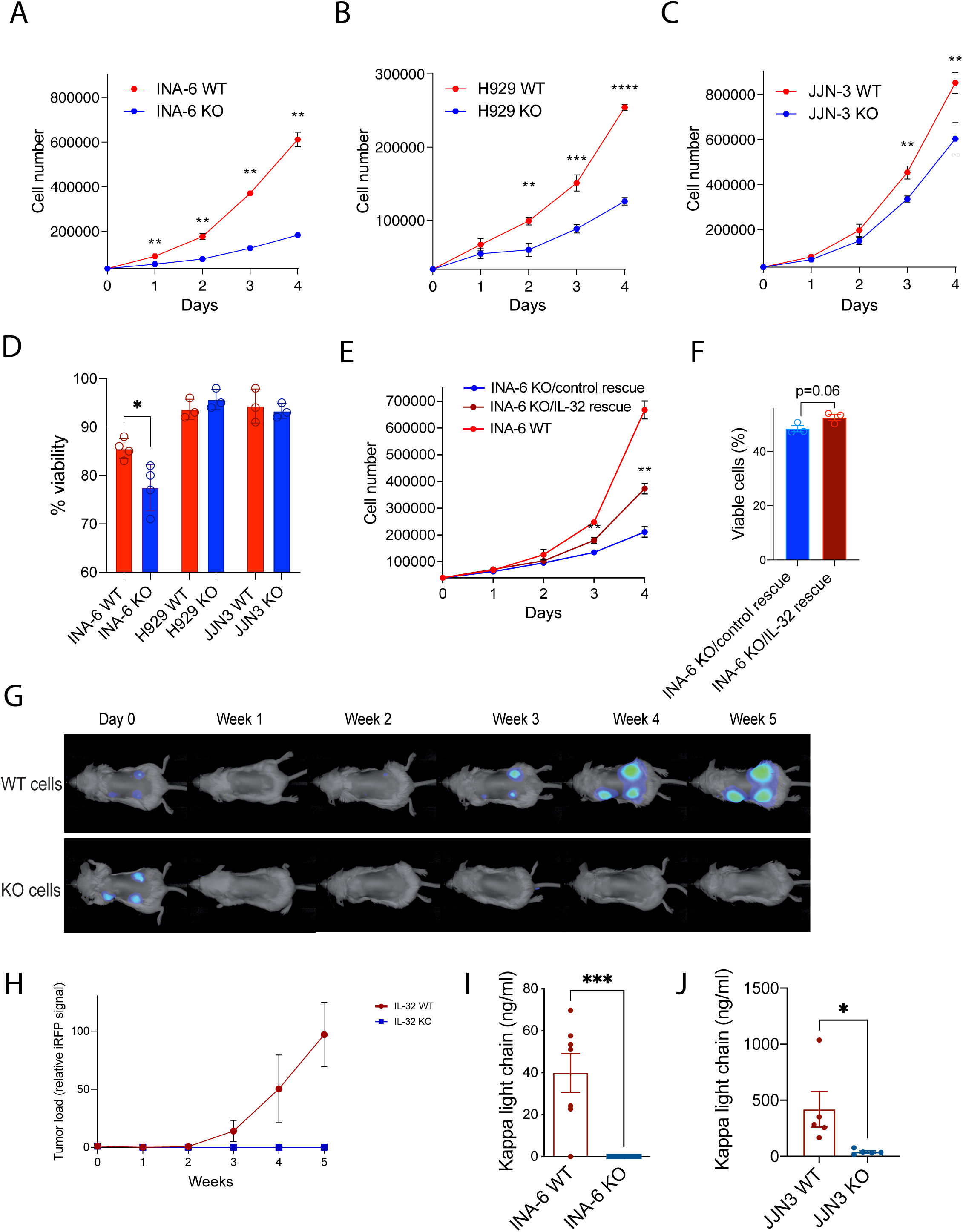
IL-32 is important for myeloma cell proliferation and survival *in vitro* and *in vivo*. **(A-C)** INA-6, H929 and JJN-3 IL-32 KO cells were generated by CRISPR/Cas9. Proliferation of IL-32 KO and WT mock cells was assessed by automated cell counting every day for 4 days. Mean ± SD of 3 technical replicates of one representative experiment of ≥3 independent experiments are shown. Significance was evaluated by calculating mean for each day and performing multiple t-tests. **(D)** Viability of INA-6, H929 and JJN-3 IL-32 KO and WT mock cells was evaluated by trypan blue staining. Data shown are mean ± SEM ≥3 independent experiments. Statistical significance was determined by unpaired Student’s t-test. **(E)** IL-32 was reintroduced into INA-6 KO cells by transduction with an IL-32 lentiviral vector and proliferation of INA-6 KO/IL-32 rescue cells and INA-6 KO/ control rescue cells was assessed by cell counting. Mean ± SD of 3 technical replicates of one representative experiment of ≥3 independent experiments are shown. Significance was evaluated by calculating mean for each day and performing multiple t-tests. **(F)** Viability of INA-6 KO/IL-32 rescue cells and INA-6 KO/control rescue was evaluated by flow cytometry using annexin/PI staining. Data shown are mean ± SEM ≥3 independent experiments. Statistical significance was determined by unpaired Student’s t-test. **(G)** 1×10^6^ iRFP labelled INA-6 IL-32 KO and WT mock cells were implanted on humanized bone scaffolds on the flanks of RAG -/- *yc*-/- BALB/c mice and tumor burden was assessed every week. The figure shows representative images of tumor burden mice injected with WT mock and KO cells. WT: N=9, KO: N=10. **(H)** Tumor development quantified by the pooled iRFP signal of all scaffolds. Figure shows mean ± SEM of WT: 27 scaffolds, KO: 30 scaffolds. **(I)** Blood was collected at the end of the experiment described in (G) and serum human kappa light chain was quantified. **(J)** 1×10^5^ JJN3 WT (N=5) or KO (N=5) cells were injected into the tibia of male RAG2−/−GC−/− mice. After 20 days blood was collected, and serum human kappa light chain was quantified. ns: not significant, * P≤ 0.05, ** P ≤ 0.01, *** P ≤ 0.001, **** P ≤ 0.0001.

IL-32 has different isoforms^13^ and based on RNA-sequencing several isoforms are expressed in the three cell lines. INA-6 cells express IL-32β and IL-32γ, with the highest expression of the β-isoform (Supplementary Figure 1C). Thus, we re-introduced IL-32β in an INA-6 KO clone (INA-6 KO/IL-32 rescue) and found that it significantly increased proliferation compared with the INA-6 KO/control rescue cells (Figure 1E) and slightly improved viability (Figure 1F), supporting that the KO phenotype was due to lack of IL-32. Expression of IL-32 in the knock-in clones was confirmed by qPCR and western blotting (Supplementary Figure 1D, E). Treating the cells with rhIL-32α, β and γ on the other hand had no effect on proliferation of the cell lines (Supplementary Figure 1F, G and data not shown) or on survival of INA-6 IL-32 KO cells (Supplementary Figure 1H). rhIL-32 was biological active since it induced TNFα production in monocytes (data not shown). Thus, intracellular IL-32, rather than exogenous IL-32 acting through a cell surface receptor, is responsible for the growth-promoting effect of IL-32 in plasma cells.

Myeloma cell growth and survival are aided by factors secreted from cells in the BM microenvironment. To address if the loss of IL-32 could be compensated by factors produced by a human bone marrow-like environment we implanted 1 × 10^6^ INA-6 iRFP-labelled IL-32 KO and WT cells into humanized bone scaffolds in immune compromised female RAG2^−/−^ GC^−/−^mice and followed tumor growth by imaging.^19, 20^ Cell injections were successful for all mice since fluorescence was detected in all scaffolds at day 0, but only cells expressing IL-32 engrafted (Figure 1G-I). We also explored if depletion of IL-32 from the more aggressive and robust cell line JJN3 was evident in vivo. To this end we injected 1 × 10^5^ JJN3 IL-32 WT or KO cells into the tibiae of male RAG2^−/−^GC^−/−^ mice. After 20 days blood was collected, and serum human kappa light chain was quantified. Human kappa light chain was detected in all mice, but it was significantly reduced in mice injected with IL-32 KO cells (Figure 1J). Hence, loss of IL-32 in the MM cells cannot be compensated by microenvironmental-derived factors.

### IL-32 is localized to the mitochondria and interacts with components of the mitochondrial respiratory chain

An IL-32 receptor is not identified, and it is not entirely clear how IL-32 acts at the molecular level^13^. Thus, to identify IL-32 binding partners we performed co-immunoprecipitation of endogenously expressed IL-32 followed by mass spectrometry analyses of the precipitates. Pulldown was performed on lysates from cells cultured for 24 hours in hypoxic conditions (2% O_2_) to increase IL-32 protein expression^15^. IL-32 KO cells were used as pulldown control to increase the specificity of the analysis. Intriguingly, 7 of 33 proteins that bound to IL-32 were mitochondrial proteins (Table 1). Considering the proportion of mitochondrial proteins in the human proteome, this is more than could be expected by chance (chi square test with yate’s correction p=0.0005). The interacting proteins included a subunit of the ATP synthase (ATP5D), a subunit of the NADH:ubiquinone oxidoreductase (NDUFA12), which is part of the respiratory complex (RC) I subunit and a subunit of RC III, ubiquinol-cytochrome c reductase (UQCR11). IL-32 also interacted with dihydroorotate dehydrogenase (DHODH), which associates with RC III in the inner mitochondrial membrane^21^. IL-32 also pulled down the mitochondrial transporters (ABCB6 and ABCB10), involved in heme synthesis and oxidative stress response^22, 23^. Interactions of IL-32 with ATP5D and NDUFA12 were verified by IP western blotting for the INA-6 (Figure 2A), IH-1 and JJN-3 cell lines (Supplementary Figure 2A, B), supporting results from the IP-MS analysis. Due to lack of suitable antibodies reverse IP with NDUFA12 and ATP5D was not possible. We were however able to pull down IL-32 using an antibody toward the ATP synthase complex (Supplementary Figure 2C), further supporting an association between IL-32 and the ATP synthase. Localization of IL-32 to the mitochondria was confirmed by the presence of IL-32 in the mitochondrial fraction of cell lysates (Figure 2B). (Supplementary Figure 2F). IL-32 was also found colocalized with mitochondria at distinct sites by confocal microscopy (Figure 2C).

**Figure 2:**
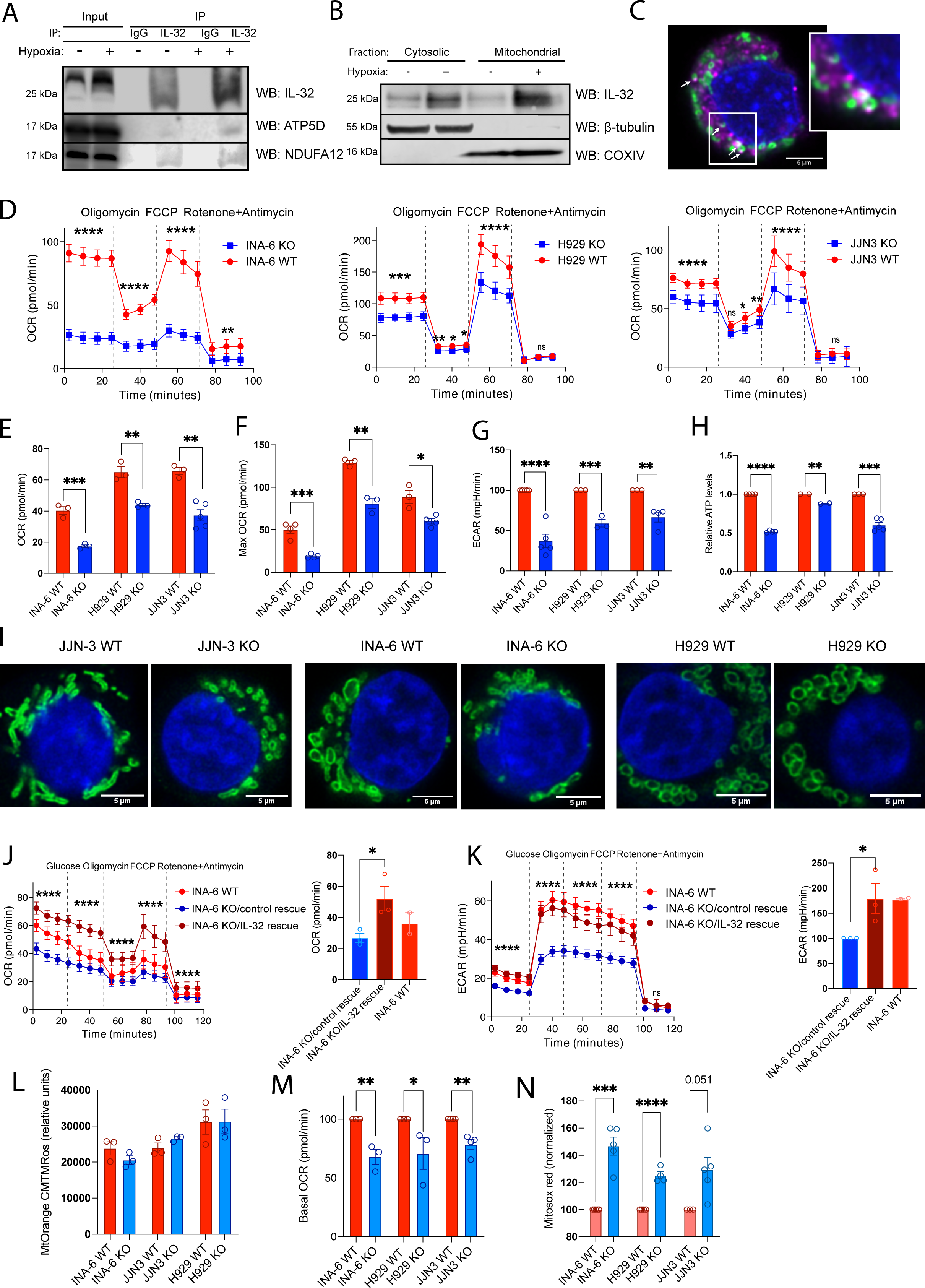
IL-32 interacts with components of the electron transport chain and enhance mitochondrial respiration. **(A)** CO-IP was performed by pulldown of endogenous IL-32 in INA-6 cells (N=3). Representative western blot of ATP5D, NDUFA12 and IL-32 is shown. **(B)** Representative western blot of IL-32 in the mitochondrial and cytosolic fraction of JJN-3 cells cultured in normoxia (20 % oxygen) and hypoxia (2 % oxygen). **(C)** Representative confocal image of hypoxic JJN-3 cells stained for IL-32 (magenta, Alexa 647), mitochondria (TOMM20, green, Alexa 488) and nucleus (blue, Hoechst). Imaging was performed with a Leica SP9, using a 63X 1.4 (oil) objective and LAS X software and deconvoluted using Huygens. Scale bar: 5μM. Arrows indicate areas of colocalization of TOMM20 and IL-32. **(D)** Representative seahorse mito stress assay measuring OCR in INA-6, H929 and JJN-3 KO and WT mock. 4 first measurements show basal OXPHOS, after injection of oligomycin: non-ATP oxygen consumption (proton leak), after FCCP injection: maximal OCR, after injection of rotenone and antimycin: non-mitochondrial respiration. Data shows mean±SD of 20 technical replicates. The differences between KO and WT mock cells were significant using Two-way ANOVA and Sidàk’s multiple comparison test (P ≤ 0.0001). **(E)** Mean basal respiration (basal OCR) in INA-6, H929 and JJN-3 KO and WT mock cell lines. Data shown is mean ±SEM of 3 independent experiments. **(F)** Mean maximal respiration (max OCR) in INA-6, H929 and JJN-3 KO and WT mock cell lines. Data shown is mean ±SEM of 3 independent experiments. **(G)** Mean basal glycolysis (±SEM) in IL-32 KO and WT cell lines analyzed by seahorse glycolysis stress test measuring ECAR. Data shown is mean ±SEM of 3 independent experiments. **(H)** Relative ATP levels in INA-6, H929 and JJN-3 KO and WT mock cells quantified by CTG-assay. Data shown is mean ±SEM of 3 independent experiments. **(I)** Representative confocal images of mitochondria of IL-32 JJN-3 KO and WT mock cells were stained for TOMM20 (green, Alexa 488) and nuclei (Hoechst, blue) and imaged as described in (C). **(J)** Representative graph showing OXPHOS in INA-6 KO/IL-32 rescue cells and IL-6 KO/rescue control (mean ± SD of more than 20 technical replicates). The difference between INA-6 control rescue and INA-6 IL-32 rescue was significant using Two-way ANOVA (P ≤ 0.0001). Bar plot shows mean basal OCR (±SEM) of 3 independent experiment. INA-6 WT mock was included for comparison. **(K)** Representative graph showing glycolysis in INA-6 IL-32 rescue cells and rescue control (mean ± SD) of more than 20 technical replicates). The difference between INA-6 control rescue and INA-6 IL-32 rescue was significant using Two-way ANOVA and Sidàk’s multiple comparison test (P ≤ 0.0001). The bar plot shows mean basal glycolysis (ECAR) (±SEM) of 3 independent experiment. INA-6 WT mock was included for comparison. **(L)** Membrane potential in isolated mitochondria from IL-32 KO and WT mock cells quantified by Mitotracker Orange CMTMRos fluorescence. The bar plots show mean ±SEM of 3 independent experiments. **(M)** Mean basal respiration (basal OCR) in isolated mitochondria from INA-6, H929 and JJN-3 KO and WT mock cell lines. Data is shown as mean ±SEM of 3 independent experiments. **(N)** Mitochondrial ROS in INA-6, H929 and JJN-3 KO and WT mock cell lines quantified by Mitosox Red staining. Figure shows Mitosox fluorescence of KO and WT cells normalized to WT for each independent experiment (N>3). * P≤ 0.05, ** P ≤ 0.01*** P ≤ 0.001, **** P ≤ 0.0001.

**Table 1:**
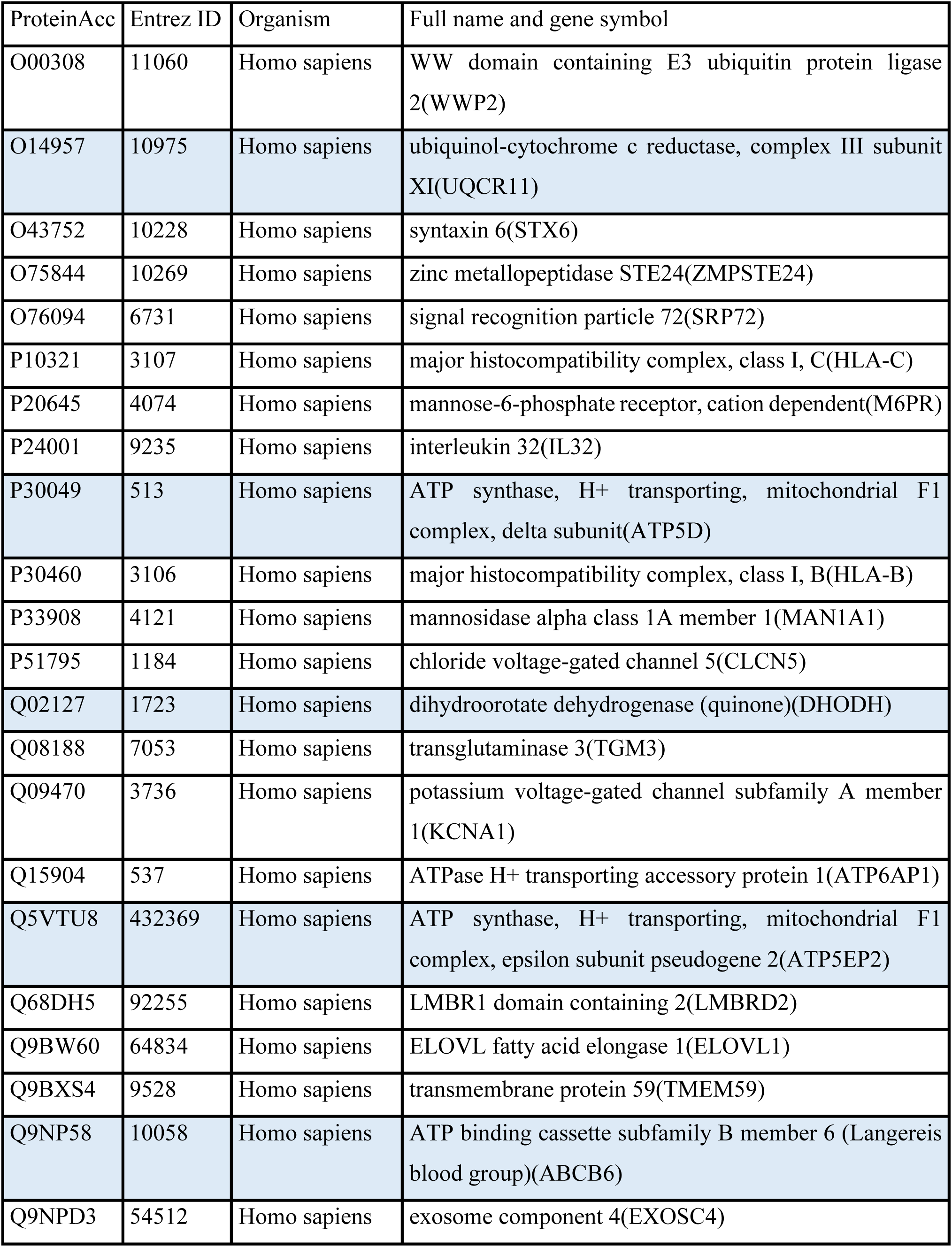
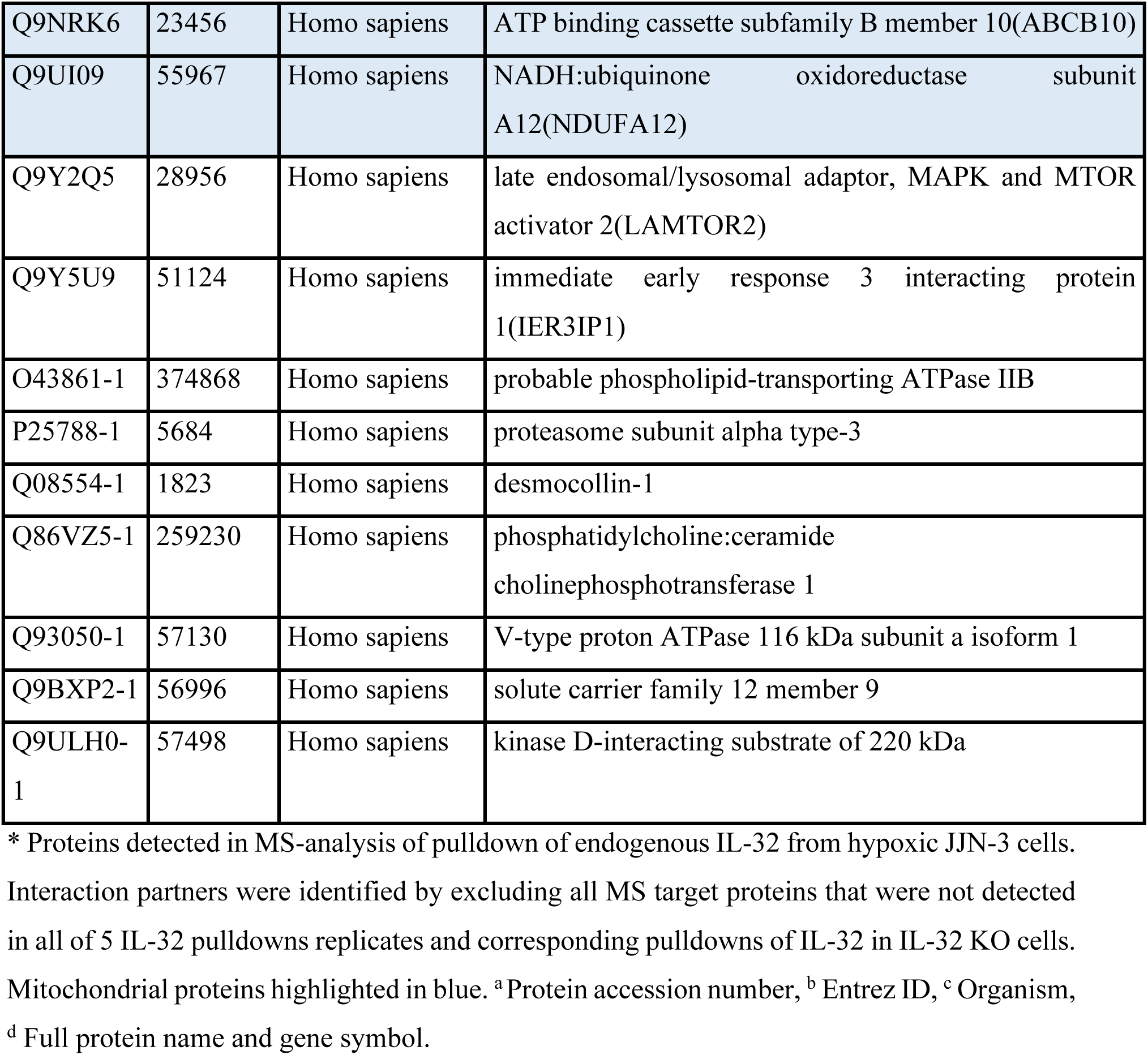
Proteins identified as interaction partners for IL-32*.

### IL-32 enhances mitochondrial respiration

To investigate if IL-32 regulates mitochondrial respiration we measured oxidative phosphorylation (OXPHOS) by quantifying the oxygen consumption rate (OCR). OCR was significantly reduced in all three IL-32 KO cell lines (Figure 2D). The IL-32-expressing cells respired significantly more than KO cells both in basal culture conditions (Figure 2E) and when maximum respiration was triggered by FCCP (Figure 2F), supporting that IL-32 promote OXPHOS in MM cells. Glycolysis, as measured by extracellular acidification rate (ECAR) was also significantly reduced in KO cells compared to WT cells (Figure 2G). Thus, aerobic glycolysis did not seem to be increased to compensate for the lack of aerobic respiration. In line with the reduction in OCR, intracellular ATP was reduced in IL-32-KO cells compared with WT cells (Figure 2H). The mitochondria in JJN-3 and INA-6 KO cells appeared rounded and small, compared with the more elongated, fused mitochondria of WT cells (Figure 2I). However, neither the amount mitochondria (Supplementary Figure 2D) nor the mitochondria membrane potential was changed (Supplementary Figure 2E). Thus, the reduction in OCR and ATP production was due to less efficient OXPHOS in the mitochondria rather than a general depolarization of mitochondria or reduced amount mitochondria in the KO cells. Transfection of IL-32β into INA-6 KO cells led to expression of IL-32β in both cytosol and mitochondria (Supplementary Figure 2F). The INA-6 KO/IL-32 rescued cells had improved metabolic capacity as both OCR and ECAR was increased (Figure 2J, K) supporting that the metabolic phenotype was due to lack of IL-32.

To further explore the effect on IL-32-depletion on mitochondrial function we isolated mitochondria from WT and KO cells. In line with the results performed on whole cells (Supplementary Figure 2D) the membrane potential did not differ in mitochondria isolated from WT and KO cells in the three cell lines (Figure 2L). Importantly, however, isolated mitochondria from KO cells showed reduced OCR (Figure 2M). These findings support that the reduced OCR was related to a reduced efficiency of mitochondrial respiration and not a result of reduced availability of TCA substrate (pyruvate) from glycolysis or other anaplerotic substrates. Despite reduced OCR (Figure 2D), a significant increase in mitochondrial ROS (mtROS) was measured in whole cells (Figure 2N). There were also more ROS in isolated mitochondria (Supplementary Figure 2G). Increased ROS may be due to electron leak in the mitochondrial electron transport chain when electrons exit prior to the reduction of oxygen to water at cytochrome c oxidase ^24^. Increased mtROS, despite of reduced OCR/ATP synthesis may thus be due to dysfunctional/suboptimal ETC.

### Loss of IL-32 leads to perturbations in metabolic pathways and downregulation of genes involved in stress response, proliferation and immune activation

Since IL-32 seemed to influence mitochondrial function, we characterized the metabolome and lipidome of INA-6 WT and KO cells from two different clones using mass spectrometry. The INA-6 cell line was chosen since it is IL-6-dependent and relatively similar to primary cells^25^. We observed major differences in metabolites between KO and WT cells (Figure 3A). There was a striking accumulation of polyunsaturated triglycerides (TAGs) in the KO cells; 36 of the 89 significant upregulated metabolites were TAGs (Figure 3B, C, Supplementary Table 1). D-fructose and citrate were also on top of the list of metabolites increased in KO cells (Figure 3B). On the other hand, 85 of 220 downregulated metabolites were membrane lipids (Supplementary Table 1), including phosphatidylethanolamines (PE), phosphatidylcholines (PC), diacylglycerols (DAG) and saturated TAGs (Figure 3C). This indicates that fatty acid synthesis is skewed in KO cells and that fatty acids are used for synthesis of unsaturated triglycerides rather than membrane lipids. Indeed, when staining for neutral lipids in JJN-3 and INA-6 KO cells we observed a striking accumulation of lipid droplets not present in the WT cells (Figure 3D, Supplementary Figure 3A). Another interesting observation was the accumulation of ceramide species in KO (Supplementary Figure 3B), including Cer (d18:1/16:0), while sphingomyelin (44:2) was downregulated in KO2 and completely absent in KO1 (Supplementary Figure 3C). According to the Metabolite Set Enrichment Analysis (MSEA) the metabolic pathways aspartate metabolism, urea cycle, purine metabolism, the citric acid cycle, PC biosynthesis, the mitochondrial electron transport chain and Warburg effect were downregulated in KO cells (Supplementary Figure 3D).

**Figure 3:**
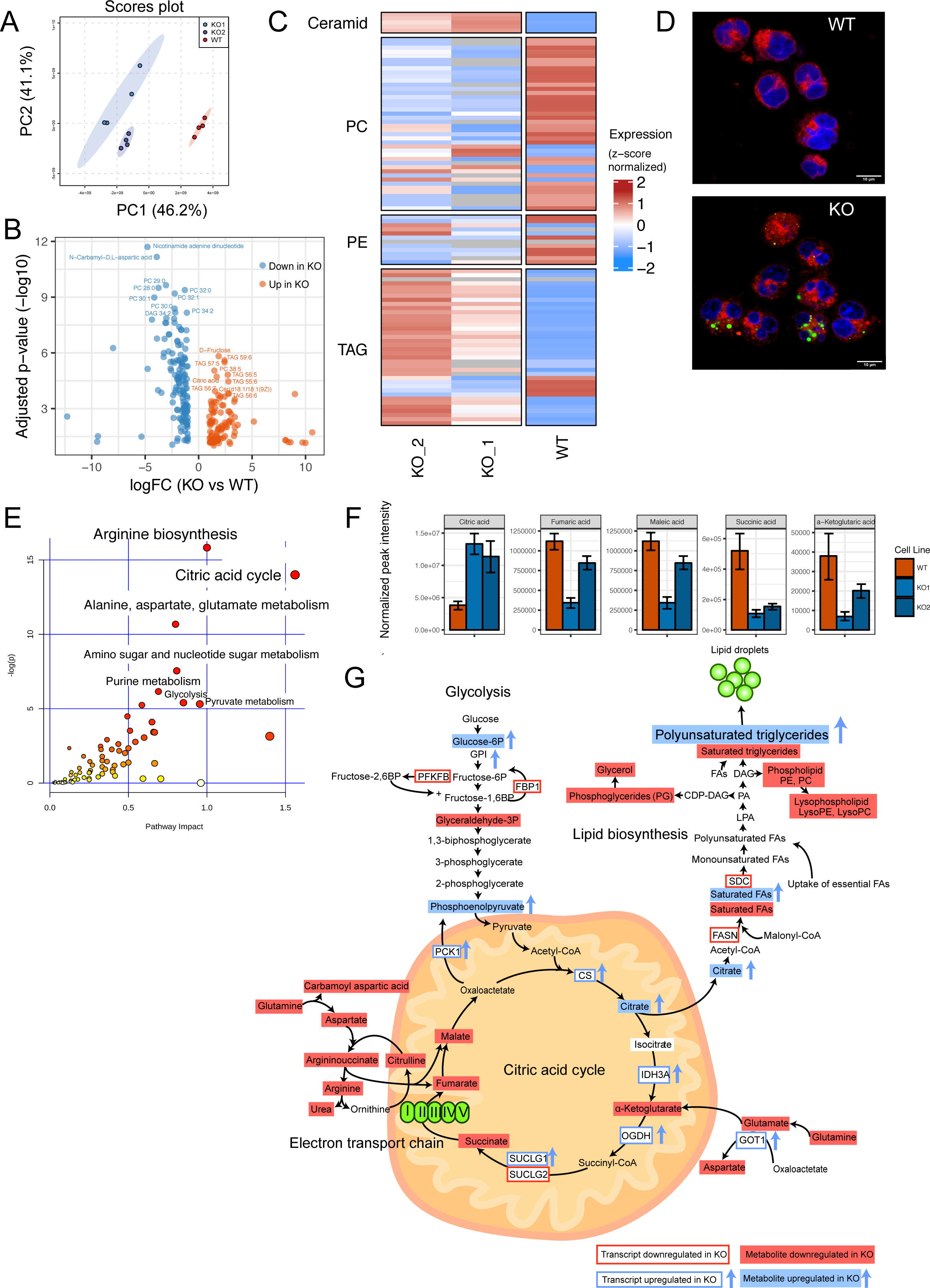
Loss of IL-32 leads to perturbations in metabolic pathways. **(A)** PCA plot of metabolomes from two clones of INA-6 KO cells and WT mock cells. **(B)** Volcano plot showing significant different metabolites (p<0.05) between KO cells and WT mock cells (metabolite expression from replicates from two KO clones were merged). **(C)** Heatmap showing significantly altered ceramides, phosphatidylcholine (PC), phosphatidylethanolamine (PE) and triglyceride (TAG) species in two INA-6 KO clones (KO1, KO2) and WT mock cells. The details on the lipid species within each class can be found in Supplementary Table 1. **(D)** Representative image of lipid droplets in INA-6 IL-32 KO and WT mock cells, stained with Nile Red. Polar lipids (red) were excited at 590 nm (600–700 nm) and neutral lipids (green) at 488 nm (500–580 nm). Confocal imaging was performed with a Leica TCS SP8 STED 3X, using a 63X 1.4 (oil) objective and LAS X software. Scale bar: 10 μM. **(E)** Joint pathway analysis (SMPDB pathways, Metaboanalyst 4.0) of transcriptomic and metabolomic data from 2 INA-6 IL-32 KO clones (KO1, KO2) and WT mock cells. **(F)** Significantly (P<0.05) altered citric acid cycle intermediates in two KO clones (KO1, KO2) vs. WT mock cells. **(G)** Illustration of significantly differentially expressed genes and metabolites from the most enriched pathways in the joint pathway enrichment analysis in G. Significance was determined by two-sided Student’s T-test using Metaboanalyst 4.0 software.

RNA sequencing of the cell lines also demonstrated marked differences in gene expression between IL-32 WT and KO cells (Supplementary Figure 4A-E, Supplementary Table 2). The common downregulated genes included genes involved biological processes such as “cotranslational protein targeting to the membrane”, “nuclear-transcribed mRNA catabolic process, nonsense-mediated decay” and “translational initiation”. In addition, terms related to immune cell proliferation, immune activation and leukocyte differentiation was downregulated in the KO cells (Supplementary Figure 4F). In contrast, biological pathways that were upregulated in INA-6 KO cells included “translational termination”, mitochondrion organization” and “regulation of G2/M transition of mitotic cell cycle”. Overall, genes related to processes in the mitochondria, cell division and protein synthesis/turnover were prominently upregulated in KO cells (Supplementary Figure 4G). Considering the phenotype of the KO cells, it is likely that some of the genes involved in these biological processes are upregulated in the KO cells as a compensatory mechanism.

By combining the metabolomics and transcriptomics data for INA-6 cells in a joint pathway analysis, we found arginine biosynthesis, citric acid cycle and alanine, aspartate and glutamate metabolism to differ the most between WT and KO cells (Figure 3E). For the individual metabolites, citrate was the only upregulated intermediate in the citric acid cycle in the KO cells, while α-ketoglutarate, succinate, fumarate and malate were all downregulated (Figure 3F, I, Supplementary Table 1), indicating that the citric acid cycle is disrupted at this point in IL-32 KO cells. Limited oxidation in the electron transport chain may lead to enhanced transport of citrate out from the mitochondria and used for synthesis of fatty acids (Figure 3G).^26^ Supporting our experimental data, ATP was reduced in the KO cells (Supplementary Table 1) and NAD was the most significantly downregulated metabolite in the KO cells (Figure 3B), indicative of less active mitochondrial metabolism in the KO cells. Taken together, changes in citric acid cycle intermediates, arginine biosynthesis and fatty acid accumulation indicate dysfunctional mitochondrial OXPHOS in IL-32 KO cells.

### IL-32 expression in primary MM cells is associated with inferior survival, cell division and oxidative phosphorylation

We have previously shown that a subgroup of 10-15 % of MM patients express IL-32 and that high IL-32 expression in patients associates with reduced progression free survival^15^ (Supplementary Figure 5A,B). To further validate IL-32 as a prognostic factor, we analyzed overall survival of IL-32-expressing patients (upper 10^th^ percentile, N= 80) and IL-32-non-expressors (lower 90^th^ percentile, N= 712) in the MMRF-CoMMpass IA13 dataset. Indeed, IL-32-expressors live significantly shorter (1005 days median survival) compared with non-expressors (median survival not reached, P= 8.9e-5) (Figure 4A). IL-32 expression also retained prognostic information when adjusting for ISS stage (Supplementary Figure 5C). Moreover, when analyzing paired diagnosis and progression samples from the same dataset, IL-32 was significantly increased upon relapse in individual patients (Figure 4B).

**Figure 4:**
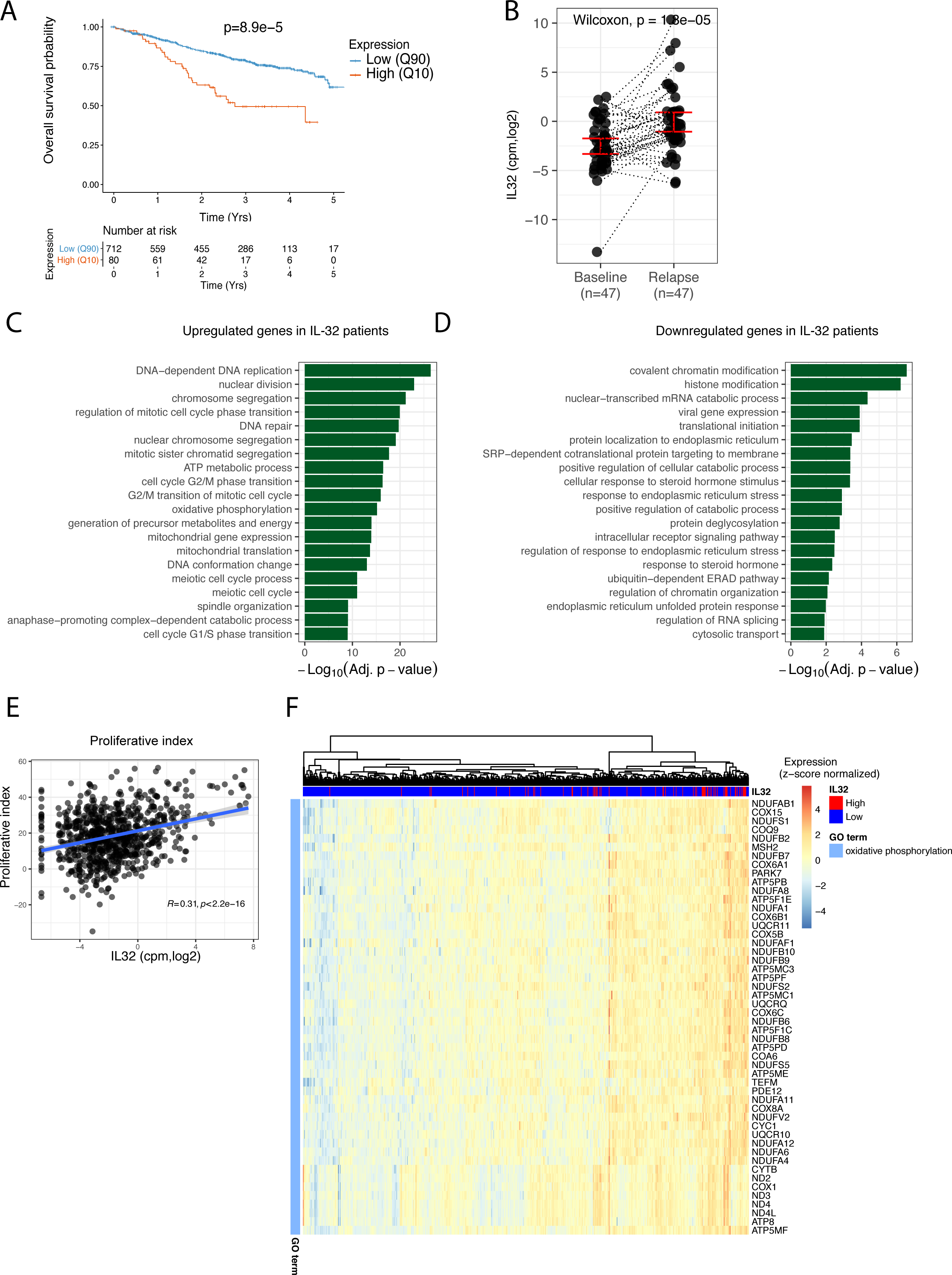
IL-32 expression in primary myeloma cells is associated with inferior survival, cell division and oxidative phosphorylation. **(A)** Overall survival of IL-32 expressing patients (10^th^ percentile) compared to non-expressing patients (90^th^ percentile) in the IA13 CoMMpass dataset P= 8.9e-5, using Cox proportional-hazards regression model. **(B)** IL-32 expression in individual patients at diagnosis and first relapse in RNA-sequenced CD138+ cells from CoMMpass IA13. Significance was determined by Wilcoxon signed-rank test. **(C)** GO-analysis of the differentially expressed genes (Benjamini-Hochberg-adjusted P-value < 0.05; log2 fold change >0 and <0 for up-and-down-regulated genes, respectively) between IL-32-expressing patients (10^th^ percentile) and IL-32 non-expressing patients (90^th^ percentile). Top significantly enriched biological processes upregulated in IL-32 expressing patients are shown. The GO terms are ordered by the Benjamini-hochberg adjusted p-values. **(D)** GO-analysis of the differentially expressed genes (Benjamini-Hochberg-adjusted P-value < 0.05; log2 fold change >0 and <0 for up-and-down-regulated genes, respectively) between IL-32-expressing patients (10^th^ percentile) and IL-32 non-expressing patients (90^th^ percentile). Top significantly enriched biological processes downregulated in IL-32 expressing patients are shown. The GO terms are ordered by the Benjamini-hochberg adjusted P-values. **(E)** Correlation between *IL32* and a proliferative index gene signature (calculated as the sum of expression values of the gene set as described in^27^). **(F)** Heatmap showing unsupervised hierarchical clustering of genes in the GO-term oxidative phosphorylation for IL-32-expressing patients (10^th^ percentile) and patients without IL-32 (90^th^ percentile).

We next examined the characteristics of IL-32-expressing primary MM cells in terms of gene expression. In the MMRF-CoMMPass IA13 dataset there were 4548 significantly differentially expressed genes between IL-32-expressing (upper 10th percentile) and non-expressing patients (lower 90th percentile) (Supplementary Table 4). Interestingly, the GO enrichment analysis of differentially expressed genes revealed changes in similar GO biological processes as associated with loss of IL-32 in the cell lines: the most **upregulated** genes in IL-32-expressing patients were associated with cell division (Figure 4C, Supplementary Table 3) indicating that this is indeed a signature of IL-32, both in cell lines and in primary cells. Moreover, IL-32 expression correlated with expression of genes associated with a high proliferative index in myeloma ^27^ (Figure 4E). OXPHOS and ATP synthesis coupled electron transport were also significantly enriched in the GO-analysis, supporting our *in vitro* data. Strikingly, of the 108 genes on the OXPHOS GO-term list, the 50 genes that were enriched in the IL-32 - expressing patients were mainly encoding subunits in the respiratory chain including subunits of the ATP synthase and subunits of the NADH:ubiquinone oxidoreductase (Figure 4F). In line with previous published data ^15^ we found IL-32 expression to be highly correlated with HIF1α expression (Supplementary Figure 6A, B). Genes **downregulated** in IL-32-expressing patients were associated with protein handling and endoplasmic reticulum stress, biological processes related to the high immunoglobulin secretion from terminally differentiated plasma cells (Figure 4D).

To investigate the distribution of IL-32 gene expression within the malignant plasma cell population and to see if the highly proliferating, respiratory phenotype is directly linked to IL-32 expression within the same cell, we analyzed a publicly available single cell dataset of MM cells sampled from bone marrow- and extramedullary tumors^28^. We identified nine distinct clusters across the 12 patient samples with a total of 488 single cells of which IL-32 was mainly expressed in three of the clusters and in four of the samples (Figure 5A-C). IL-32 was expressed in about 70% of the cells from sample MM33 and at intermediate levels in the majority of cells from MM17 as well as in a few cells from MM36 (Figure 5B-C). Interestingly, in patient MM02 IL-32 was not expressed in the bone marrow sample taken at diagnosis (MM02), but highly expressed in all the cells of the extramedullary tumor sample (MM02EM) obtained 18 months later, linking IL-32 expression to MM cells with a metastatic potential. Importantly, genes involved in “ATP synthesis coupled electron transport”, “assembly of ETC complexes”, and “cell cycle progression” were significantly upregulated in single cells expressing IL-32 compared with non-expressing cells (Figure 5D). These data support that the same MM cell that express IL-32 has high OXPHOS and proliferation.

**Figure 5:**
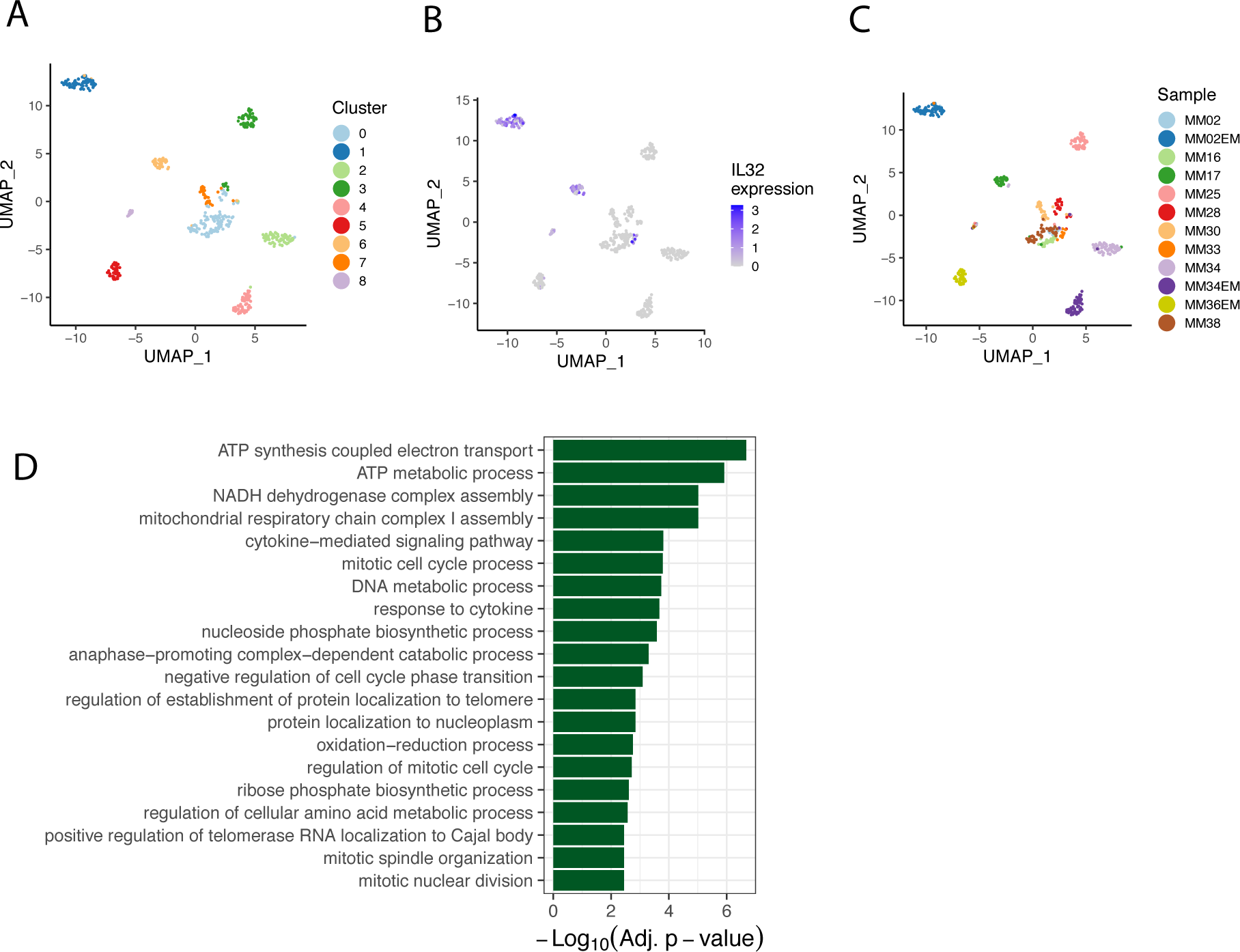
Single cell transcriptome analysis of IL-32-expressing myeloma cells. **A)** UMAP plot colored by the identified clusters. **B)** UMAP plot colored by the level of IL32-expression per cell. **C)** UMAP plot colored by patient sample. **D)** Top 20 gene ontology terms (biological processes) for genes enriched in IL-32 expressing patient cells. The GO terms are ordered by the Benjamini-hochberg adjusted p-values. The data were obtained from Ryu et al. ^28^.

### IL-32 expression promotes a more immature plasma cell phenotype

Changes in metabolism and metabolites may change gene expression at several levels^29^. To gain further knowledge of the transcriptional programs associated with IL-32 in malignant plasma cells, we investigated which genes were downregulated in the INA-6 KO cells and at the same time upregulated in IL-32-expressing primary cells in CoMMpasss IA13 dataset. (Supplementary Table 4, Figure 6A). We identified 230 genes to be significantly differently expressed in both comparisons, and these genes are likely to be functionally related to IL-32 expression. The top 3 genes, when sorting for the most downregulated genes in KO and upregulated genes in IL-32-expressing patients on the shared signature list was *MME*, encoding the receptor CD10, the transcription factor *IRF8* and *SORL1*, encoding the sortilin-related receptor 1 (Figure 6B, C). SORL1 plays a role in lipid metabolism and IL-6 signaling ^30–32^ and *MME* and *IRF8* are both important in early stages of B-cell development ^33, 34^. *MME* and *SORL1* were also downregulated in H929 KO cells compared to WT cells (Supplementary Figure 7A). *IRF8* was not expressed by this cell line.

**Figure 6:**
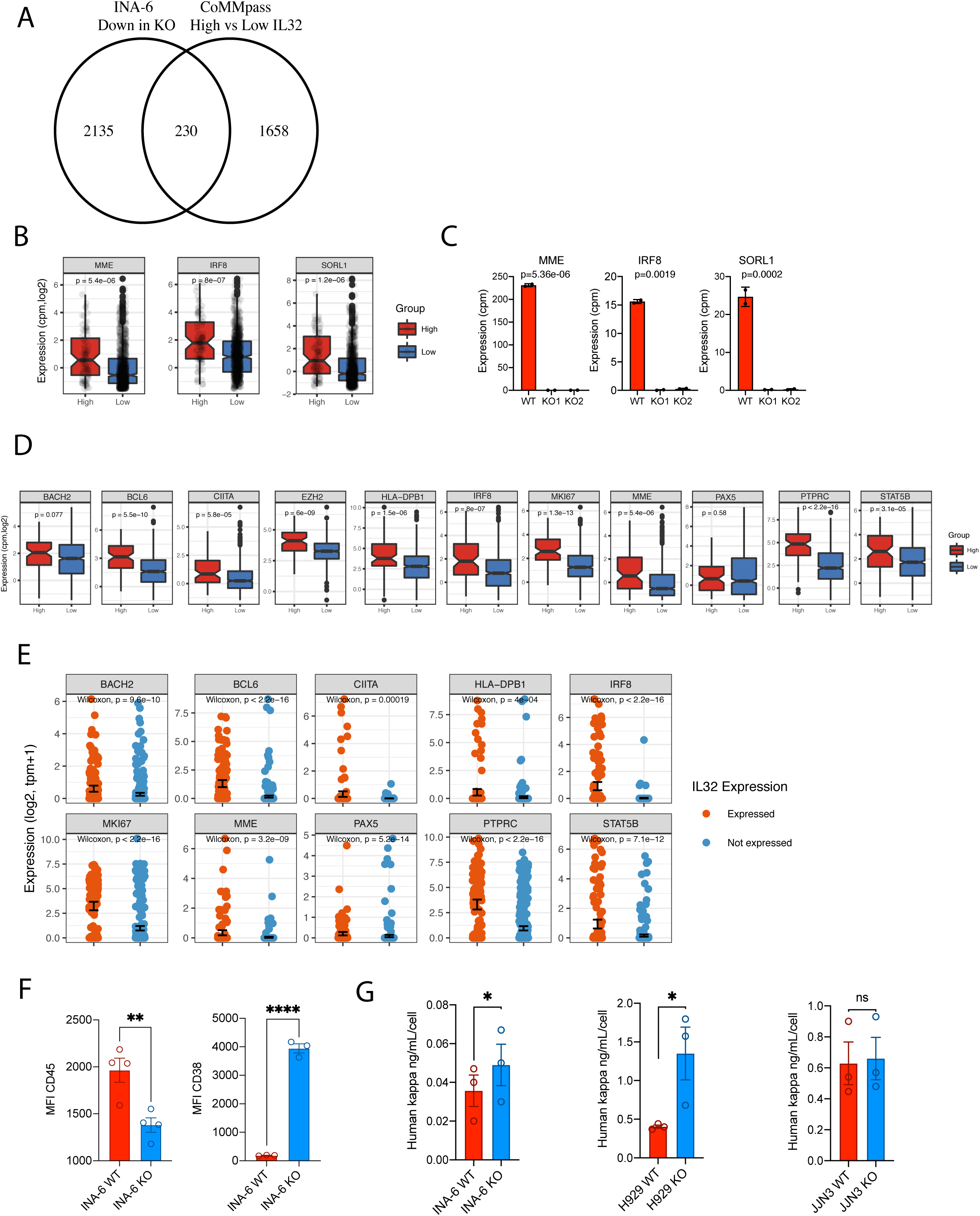
IL-32 expression promotes a more immature plasma cell phenotype. (**A**) Venn-diagram of overlapping significant genes (p>0.01) that were downregulated in KOs (comparing two INA-6 KO clones (KO1, KO2) with WT mock cells) and upregulated in IL-32 patients (comparing IL-32-expressing patients vs non-expressing patients). (**B**) Gene expression of *MME*, *IRF8* and *SORL1* in patients expressing IL-32 (10^th^ percentile) compared to non-expressing (90^th^ percentile) patients. Significance determined by Wilcoxon signed-rank test in *R*. (**C**) Gene expression of *MME*, *IRF8* and *SORL1* in INA-6 IL-32 KO1, KO2 and WT mock cells. Significance determined by limma in *R* with Benjamini-Hochberg-adjusted P-values. (**D**) Evaluation of gene expression of markers associated with less differentiated stages of B-cell maturation in CoMMpass IA13, comparing IL-32 expressing patients (upper 10^th^ percentile) to non-expressing patients (lower 90^th^ percentile). Significance determined by Wilcoxon signed-rank test in *R*. (**E**) Scatterplot of same genes as in (D) in single cells with (N=142) and without (N=346) IL32-expression (from single cell transcriptomics). P-values were calculated from a two-sided Wilcoxon signed-rank test in *R*. (**F**) Surface expression of CD45 and CD38 in INA-6 KO and WT cells. Data is presented as median fluorescence intensity (MFI) from 3 independent experiments and significance determined by unpaired student’s T-test. (**G**) Concentration of kappa light chain/cell detected in conditioned media from WT and KO cells as indicated. P-values were calculated by the ratio paired t-test.

Other genes associated with earlier stages of B cell differentiation^35–40^ such as *BCL6*, *BACH2*, *EZH2*, *STAT5B*, *PTPRC* (CD45), *MKI67* and several genes encoding MHC II, including *HLA-DPB1* were upregulated in IL-32-expressing patients (Figure 6D, Supplementary Table 4). Importantly, we also found expression of the same genes to be significantly upregulated in IL-32-expressing cells in the single cell sequencing dataset (Figure 6E). In accordance, genes associated with mature plasma cells were slightly, but significantly downregulated in IL-32-expressing patients, including *CD27*, *CXCR4*, *ERN1* and *FOXO1* (Supplementary Figure 7B). CD45 is as marker of proliferating, immature myeloma cells, while CD38 is a marker of terminally differentiated plasma cells and expressed by the majority of MM cells ^41^. INA-6 is a MM cell line with an immature phenotype, with high expression of CD45 and relatively low expression of CD38. Strikingly, loss of IL-32 led to a reduction in CD45 and re-expression of CD38 in INA-6 cells as examined by flow cytometry (Figure 6F). Other genes associated with immature plasma cells were also downregulated in the KO cells (Supplementary Figure 7C). ER stress is an Achilles heel of MM cells partly due to the production of large amounts of monoclonal antibodies. IL-32-expressing primary cells seemed less affected by ER stress (Figure 4D), so we asked whether this could be related to less production of immunoglobulins. Strikingly, INA-6 and H929 KO cells secreted more kappa light chain than WT mock cells (Figure 6G), supporting our hypothesis that IL-32 expression promotes an immature plasma cell phenotype, and that IL-32-expression may relieve the cells from immunoglobulin-related ER stress. Taken together these results suggests that IL-32 is involved in regulation of transcriptional programs that induce a more immature and less ER-stressed plasma cell.

## Discussion

We have identified IL-32 as a novel, endogenously expressed growth and survival factor for malignant plasma cells. IL-32 interacts with components of the respiratory chain, and expression of IL-32 is important for efficient OXPHOS in MM cells. A subgroup of MM patients expresses IL-32, and these patients have reduced OS. Furthermore, the malignant plasma cells of these patients have distinct phenotypical characteristics, resembling an immature or less differentiated plasma cell.

Based on gene expression data in CoMMpass, 10-15% of MM patients express IL-32 at diagnosis. Moreover, analyses of paired samples from diagnosis and relapse in individual patients suggest that IL-32 expression increases upon relapse in about 20% of the patients. The strong, independent association of IL-32 with inferior survival in patients, the reduced proliferative rate and the reduction of OXPHOS in three phenotypically different cell lines when depleting IL-32 strongly suggest that IL-32 plays a role in MM disease progression. The IL-6-dependent cell line INA-6 was in particular dependent on IL-32 expression, since loss of IL-32 not only reduced proliferation but also reduced cell survival. In fact, these cells were not able to form tumors *in vivo*, even in a supportive, humanized bone marrow microenvironment. This can possibly be explained by IL-32-depleted cells being less able to adapt to the more challenging metabolic conditions *in vivo*, where there may be limited access to oxygen and changed composition of nutrients^42, 43^.

In contrast to the majority of MM cells, which in general are slow growing and have low Ki-67,^44, 45^ IL-32-expressing primary MM cells have a gene signature related to cell division and an immature plasma cell phenotype. This was evident both from the large CoMMPass dataset and from the small number of single cell RNA-sequenced patient samples. And strikingly, IL-32-depleted MM cells had downregulated expression of the same “immature” genes compared with WT cells. Metabolites have a central role in regulating gene expression. For example, availability of acetyl-CoA can modify extent of histone acetylation, while metabolites such as succinate and a-ketoglutarate may regulate DNA- and histone methylation^29^. Although additional experiments are needed to conclude, it could be that IL-32 expression and subsequent changes in metabolism leads to plasma cell de-differentiation or maturation arrest. The profound changes in ATP and core metabolites such as citrate, a-ketoglutarate, succinate, fumarate and malate upon IL-32 depletion as shown here makes this a likely scenario. Supporting this notion, GO-terms related to histone-modifications were also differently expressed between IL-32-expressing and non-expressing primary cells. Of note, MM cells with an immature phenotype have previously been linked with more aggressive tumors^41, 46^ and compounds such as ATRA and 2-methoxyestrodiol (2-ME2) that promote differentiation of MM cells render the cells more sensitive to bortezomib^47^.

Oxygen is a key regulator of aerobic respiration and metabolism and it is striking that IL-32 expression is regulated by two different oxygen sensing systems, HIF1α^15^ and cysteamine (2-aminoethanethiol) dioxygenase (ADO)^16^. ADO is an enzymatic O_2_ sensor, and was shown to catalyze dioxygenation of IL-32 in the presence of O_2_, leading to proteasomal degradation ^16^. Correspondingly, hypoxia leads to stabilization of HIF1α, which induces IL-32 mRNA and protein expression^15^. These data support that IL-32 has an important role in cellular responses to alterations in oxygen levels. Hypoxia is known to cause changes in the composition of ETC complexes and the changes help keep the mitochondria intact under low oxygen conditions and to prevent excessive ROS formation^48^. Indeed, we found that 7 of 33 proteins that bound to IL-32 in hypoxia were located in the mitochondrion, and that 5 of these were subunits of different components of the mitochondrial respiratory chain. Respirasome supercomplexes, where the respiratory chain components are assembled in close vicinity to each other, lead to higher catalytic activity of the individual components, to increased efficiency of electron transfer and to less production of ROS^24, 49^. The IL-32 KO cells had reduced capacity for mitochondrial respiration and ATP formation, still, cells lacking IL-32 had significantly higher levels of mitochondrial ROS, in line with suboptimal respiration. Thus, we propose that IL-32 by binding components of the respiratory chain enhance the efficiency of the ETC, enabling the cells to maintain OXPHOS even under conditions of low O_2_ and also to keep mtROS at a level compatible with cell survival. Exactly how IL-32 acts in the mitochondria to enhance OXPHOS needs to be further explored.

In this study we show that IL-32 is expressed by a fraction of MM cells and that it promotes survival and proliferation of these cells. IL-32 is also expressed by regulatory or senescent/exhausted T cells in MM, but if/how this affects the function of the T-cells is not known^50, 51^. IL-32 is secreted from the MM cells and IL-32 in the BM microenvironment enhance osteoclast differentiation^15^, it can promote IL-6 production from stromal cells^52^ and may generate immunosuppressive macrophages, ^53^ all factors that may lead to a more aggressive disease development.

In conclusion, we have shown that intracellular IL-32 promotes OXPHOS and provides a survival benefit for malignant plasma cells. The direct interaction of IL-32 with components of the respiratory chain and its regulation by two different oxygen sensing system indicate that IL-32 has an important role in cellular responses to O_2_ fluctuations. Besides identifying IL-32 as a potential prognostic biomarker and treatment target in MM, our results provide novel insight into the metabolic functions of IL-32, which may be further exploited in other cancers and inflammatory diseases where IL-32 is known to play a central role.

## Methods

### Cells and culture conditions

MM cell lines INA-6 and JJN-3 were kind gifts from Dr. Martin Gramatzki (University of Erlangen-Nuremberg, Erlangen, Germany), and Dr. Jennifer Ball (University of Birmingham, UK), respectively. H929 cells were obtained from ATCC. H929 and JJN-3 were cultured in 10% heat inactivated fetal calf serum (FCS) in RPMI-1640 (RPMI) medium. INA-6 cells were cultured in 10% FCS in RPMI with the addition of 1 ng/ml recombinant human (rh) interleukin (IL)-6. Cell lines were maintained at 37°C in a humidified atmosphere with 5% CO_2_.

### Generation of IL-32-depleted cells

IL-32 KO cell lines was generated by lentiviral transduction or plasmid transfection. For lentiviral transduction of NCI-H929 cells, the IL-32 KO CRISPR/cas9 lentiviral vector pLV-U6-gRNA/EF1a-puro-2A-CAs9-2A-GFP. # HS0000050421 and control lentiviral particles (U6-gRNA/EF1a-puro-2A-Cas9-2A-GFP) were purchased from Merck/Sigma. The CRISPR IL-32 KO vector was packaged into viral particles in HEK293T cells (Open Biosystems, Thermo Fisher Scientific), using second generation packaging plasmids psPAX2, and pMD2.G. After 48 hours, virus particles in supernatants were transduced into the target cells incubated with polybrene. Ready to use control virus particles were delivered to the cells by spin transduction. The cells were then subjected to negative selection using puromycin followed by single cell cloning. JJN-3 and INA-6 cells were electroporated with IL-32 CRISPR/Cas9 plasmid containing green fluorescent protein (GFP) for selection (#sc-406489), using buffer R (Amaxa Nucleofector Kit R VCA-1003; Lonza) and program R-001. After 24 hours, the cells were sorted for GFP positivity by fluorescence-activated cell sorting and cloned. Clones were screened for IL-32 expression by flow cytometry and immunoblotting. For all cell lines: clones that had been transfected with the plasmid (GFP^+^), but still expressed IL-32 were used as control cells (IL-32 wild type (WT)). IL-32 protein knockout was regularly confirmed by immunoblotting to ensure homogeneity of the KO cell lines.

### Generation of IL-32 knock-in cells

INA-6 IL-32 KO cells were lentiviral transduced using OmicsLink™ ORF lentiviral expression system from GeneCopoeia. Vector for knock in of IL-32β: (transcript variant 8) EX-M0733-Lv121 and control vector: EX-EGFP-Lv121 were purchased from GeneCopoeia. Lentiviral packaging and transduction of cells were performed as described above and transduced cells were subjected to negative selection using puromycin.

### Generation of iRFP-labelled cells

iRFP labelling of INA-6 KO and WT cells was performed by lentiviral transduction. piRFP (Addgene) and pENTR4 (Invitrogen) were cut with SalI and NotI (Fermentas) and ligated using a T4 DNA ligase (Fermentas) to achieve a gateway compatible iRFP entry clone. The iRFP-ENTR4 was then recombinated into the pLenti-CMV-Puro-Dest by a LR (Invitrogen) gateway reaction as previously described ^20^. iRFP lentivirus was made by transfecting HEK293T using iRFP plasmid, psPAX2 and pMD2.G packaging plasmids. iRFP-positive cells were sorted on a FACSAria™ Fusion flow cytometer (BD Biosciences, San Jose, CA, USA) giving a pure iRFP-labelled population for *in vivo* studies.

### Recombinant proteins and antibodies

For rhIL-32 stimulation experiments recombinant human IL-32α (#3040-IL-050), IL-32β (#6769-IL-025) and IL-32γ (#490-IL-025) were used. The following antibodies were used: anti-IL-32 (#AF3040) from R&D Systems, anti-β-tubulin (#2128) and anti-COXIV (#4850) antibodies from Cell Signaling Technology, anti-TOMM20 (#HPA011562), anti-NDUFA12 (#HPA039903), and anti-β-actin (#A2066) antibodies from Sigma, glyceraldehyde-3-phosphate dehydrogenase (GAPDH; #Ab8245), anti-ATP5D (#ab97491) and anti-ATP synthase immunocapture antibody (ab#109867) were from Abcam.

### Assessment of proliferation and survival

Proliferation of IL-32 KO and WT mock cells was assessed by cell counting every day for 4 days using cell coulter (Beckman Coulter Diagnostics). Cell viability was assessed by Trypan blue staining or by annexin/PI flow cytometry and analyzed by FACS Diva software (BD Biosciences).

### In *vivo* experiments

RAG2^−/−^ γC^−/−^ BALB/c mice were kept in a specific pathogen free (SPF) unit, housing 3-5 mice/ IVC-cage and supplied with bedding material, nesting material and enrichment objects. Mice were given sterile food and water ad libitum. The mice were kept in a room maintaining 21–22°C, 55% humidity and 12 hours light/dark cycles including 1 hour dusk/dawn. For the INA-6 in vivo experiment 20 female mice (12-16 weeks old) were used. Human bone marrow-derived mesenchymal stromal cells (hMSC) from healthy donors were seeded on biphasic calcium phosphate (BCP) scaffolds and differentiated toward osteoblasts for 1 week *in vitro* before implanted subcutaneously on the back of the mice (4 scaffold on each mice) and left for 2 months for establishment of a differentiated human bone cell microenvironment as described previously ^19^. 3 scaffolds on each mouse were injected with 10^6 iRFP labelled INA-6 KO or WT cells and imaging was performed after injection (timepoint 0) and then every week for 5 weeks. Images were acquired at 700 nm using Pearl Impulse imaging system and data was analyzed with Image Studio Software (both from Licor Biosciences). iRFP signal from each scaffold on each timepoint was normalized to the signal at day 0. For the JJN3 in vivo experiment JJN3 1 × 10^5^ IL-32 KO (N = 5) or IL-32 WT (N = 5) cells in 10 µL of PBS were injected into the tibiae of 9- to 12-week-old RAG2−/−γc−/− mice. After 20 days, the mice were euthanized. Blood was collected for quantification of human immunoglobulin kappa light chain by ELISA (Bethyl Laboratories). Animal handling and procedures were approved by the Norwegian food safety authority (FOTS10517).

### Real-time quantitative PCR

Total RNA was isolated using RNeasy kit (Qiagen). Complementary DNA (cDNA) was synthesized from total RNA using High-Capacity RNA-to-cDNA kit (Applied Biosystems). PCR was performed using StepOne Real-Time PCR System and Taqman Gene Expression Assays (Applied Biosystems) using standard settings (2′ 50°C, 10 ′ 95°C, 40 cycles at 95°C for 15 sec, 1′ 60°C). Relative gene expression was analyzed by the comparative Ct method. Probes were as follows: human IL-32 (Hs00992441_m1) housekeeping gene TATA-binding protein (*TBP*; Hs00427620_m1) or *GAPDH*(Hs99999905_m1).

### Immunoblotting

Cells were lysed in lysis buffer (50 mM Tris–HCl, 1% NP40, 150 mM NaCl, 10% glycerol, 1 mM Na_3_VO_4_, 50 mM NaF and Complete protease inhibitor (Roche Diagnostics) and the lysates were denatured in 1× NuPage LDS sample buffer supplemented with 0,1 mM DTT for 10 min at 70°C before they were separated on 10% Bis-Tris polyacrylamide gel. Proteins were transferred to a nitrocellulose membrane using the iBlot Dry Blotting System (Invitrogen). Membranes were blocked using 5% bovine serum albumin (Sigma–Aldrich) in Tris-buffered saline with 0,01% Tween followed by overnight incubation with the primary antibodies. Detection was performed using horseradish peroxidase (HRP) conjugated antibodies (DAKO) and developed with Super Signal West Femto Maximum Sensitivity Substrate (Thermo Scientific). Images were obtained with LI-COR Odyssey Fc and analyzed using Image Studio Software (LI-COR).

### Co-immunoprecipitation and mass spectrometry

IL-32 antibody (RnD) was conjugated to M-270 beads according to the manufacturer’s instructions, using Dynabeads™ Antibody Coupling Kit and 10 mg beads on 500μL (100ug) resuspended IL-32 antibody. For CO-IP MS analysis JJN-3 KO and WT cells were incubated in hypoxia for 24 hours before lysis with 4x pellet volume RIPA buffer (1% CHAPS, 50 mM Tris, 150 mM NaCl, Complete protease inhibitor (Roche Diagnostics) and phosphatase inhibitor cocktails 2 and 3 (Sigma-Aldrich), for 1 hour at 4°C on rotation. Lysate was pre-cleared by adding 40μL beads to ∼400μL lysate for 1h at 4 degrees on rotation, before incubated with IL-32 antibody conjugated beads for 2 hours at 4°C. After washing 4 times with PBS repelleted beads were subjected to MS-digestion protocol. Beads were resuspended 150 μl 50 mM NH4HCO3, followed by addition of 7.5 μl 200 mM DTT, 55°C for 30 min. Samples were cooled to RT before addition of 15 μl 200mM IAA and incubation in RT for 30 min in the dark. Then samples were treated 1,5 μg trypsin (MS-grade) at 37°C over-night, before beads were removed and liquid sample was dried using Speedvac (Thermo Fischer Scientific).

After tryptic digestion peptides were desalted using STAGETIP as previously described ^54^. After desalting, peptides were dried down in a SpeedVac centrifuge and resuspended in 0.1% formic acid. The peptides were analyzed on a LC-MS/MS platform consisting of an Easy-nLC 1200 UHPLC system (Thermo Fisher Scientific) interfaced with an QExactive HF orbitrap mass spectrometer (Thermo Fisher Scientific) via a nanospray ESI ion source (Proxeon). Peptides were injected into a C-18 trap column (Acclaim PepMap100, 75 μm i. d. x 2 cm, C18, 3 μm, 100 Å, Thermo Fisher Scientific) and further separated on a C-18 analytical column (Acclaim PepMap100, 75 μm i. d. x 50 cm, C18, 2 μm, 100 Å, Thermo Fisher Scientific) using a multistep gradient with buffer A (0.1% formic acid) and buffer B (80% CH3CN, 0.1% formic acid): From 2-10% B in 10 min, 10-50% B in 130 min, 50-100% B in 20 min and 20 min with 100% buffer B. The HPLC were re-equilibrated with 2% buffer B before next injection. The flow rate was 250 nl/min. Peptides eluted were analyzed on QExactive HF mass spectrometer operating in positive ion- and data dependent acquisition mode using the following parameters: Electrospray voltage 1.9 kV, HCD fragmentation with normalized collision energy 32, automatic gain control target value of 3E6 for Orbitrap MS and 1E5 for MS/MS scans. Each MS scan (m/z 300–1600) was acquired at a resolution of 12000 FWHM, followed by 15 MS/MS scans triggered for AGC targets above 2E3, at a maximum ion injection time of 50 ms for MS and 100 ms for MS/MS scans.

5 replicates each of KO and WT cells were used for mass-spectrometry analysis. The IL32-specific proteins were detected by subtracting the peptides detected in the KO-cells from the peptides detected in the WT-cells. We required the peptides to be detected in all 5 WT replicates. To assess the probability of detecting a frequency of 7 mitochondrial out of 36 IP target proteins we based the expected frequency of 1100 mitochondrial proteins ^55^ and 20 000 proteins as the total number of proteins in the human proteome.

For validation of interaction partners in the mitochondrial electron transport chain we chose two candidates for validation, ATP synthase subunit delta (ATP5D) a subunit in the ATP-synthase complex (complex IV) and NADH dehydrogenase [ubiquinone] 1 alpha subcomplex subunit 12 (NDUFA12), a subunit of the NADPH dehydrogenase (complex I). IL-32 was pulled down following the same protocol as earlier, in INA-6, IH-1 and JJN-3 cells, and ATP5D and NDUFA12 were detected by western blotting.

### Metabolomics

Targeted metabolomics (GC-MS and LS-MS) of INA-6 WT and two KO cell lines were performed by MetaSysX (Potsdam-Golm, Germany). The sample preparation was performed according to the company’s standard procedure, a modified protocol from Salem et al ^56^. 10^8 cells/ replicate was used for metabolite extraction.

LC-MS Measurements (Hydrophilic and Lipophilic Analytes) were performed using Waters ACQUITY Reversed Phase Ultra Performance Liquid Chromatography (RP-UPLC) coupled to a Thermo-Fisher Exactive mass spectrometer. C8 and C18 columns were used for the lipophilic and the hydrophilic measurements, respectively. Chromatograms were recorded in Full Scan MS mode (Mass Range [100-1500]). All mass spectra were acquired in positive and negative ionization modes. Extraction of the LC-MS data was accomplished with the software REFINER MS® 11.1 (GeneData, http://www.genedata.com). Alignment and filtration of the LC-MS data were completed using metaSysX in-house software. After extraction from the chromatograms, the data was processed, aligned and filtered for redundant peaks. The alignment of the extracted data from each chromatogram was performed according to the criteria that a feature had to be present in at least 3 out of 4 replicates from one group. At this stage, the average RT and *m/z* values was given to the features. The alignment was performed for each type of measurement independently, followed by the application of various filters in order to refine the dataset, which included the removal of isotopic peaks, in-source fragments of the analytes (due to the ionization method), and redundant peaks like additional less intense adducts of the same analyte and redundant derivatives, to guarantee the quality of the data for further statistical analyses. The in-house metaSysX annotation database of chemical compounds was used to match features detected in the LC-MS polar and lipophilic platform. The annotation of the content of the sample was performed by database query of mass-to-charge ratio and the retention time of detected features within certain criteria

The metaSysX in-house database contains mass-to-charge ratio and retention time information of 7500 reference compounds available as pure compounds and measured in the same chromatographic and spectrometric conditions as the measured samples. In addition, 1500 lipids and sugar esters were putatively annotated based on the precursor m/z, fragmentation spectrum and elution patterns. The matching criteria for the polar and non-polar platforms were 5 ppm and 0.085 minutes’ _deviation from the reference compounds mass-to-charge ratio and retention time, respectively. Coeluting compounds with the same mass-to-charge ratio were all kept and the names are separated with “|”. Lipid annotation was additionally performed and confirmed by MS/MS fragmentation spectrum using the metaSysX developed-R-based algorithm. This information is combined with the information from the annotation after the query of the MSX database.

GC-MS measurements were performed on an *Agilent Technologies* GC coupled to a *Leco Pegasus HT* mass spectrometer which consists of an EI ionization source and a TOF mass analyzer (column: 30 meters DB35; Starting temp: 85°C for 2min; Gradient: 15°C per min up to 360°C). NetCDF files that were exported from the Leco Pegasus software were imported to “R”. The Bioconductor package TargetSearch ^57^ was used to transform retention time to retention index (RI), to align the chromatograms, to extract the peaks, and to annotate them by comparing the spectra and the RI to the Fiehn Library and to a user created library. The annotation of peaks was manually confirmed in Leco Pegasus. Analytes were quantified using a unique mass. Metabolites with an RT and a mass spectra that did not result in a match in the database were kept as not assigned metabolites. Statistical analysis of significantly upregulated and downregulated metabolites in INA-6 KO clones compared to WT, metabolite enrichment, and joint pathway analysis for metabolomics and transcriptomics (RNA-seq) data was performed using Metaboanalyst 4.0 software.

### Confocal imaging

For IL-32/mitochondria colocalization studies cells were seeded in poly-L-lysine coated 96 well glass bottom plates and left to attach for 20 min at 37°C before fixed with 4% PFA, 10 min at RT. After quenching for 10 min with 50 mM NH_4_Cl, permeabilization was performed using 0.05% saponin in PBS. Primary antibody cocktail (2 µg/mL) diluted in 1% HS, 0.05% saponin was left on overnight at 4°C. Next day, secondary antibody (2 µg/mL) in 1% HS, 0.05% saponin was added for 30 min, before leaving cells in Hoechst (Thermo Fisher) in PBS (2 µg/mL) for imaging. For lipid droplet staining with Nile Red cells were attached to poly-L-lysine coated 96-well glass bottom plates, before fixed with 4% PFA, 10 min at RT and stained with 500 nM Nile Red (Thermo Fisher) for 10 min at 37°C. Polar lipids were excited at 590 nm (600–700 nm) and neutral lipids at 488 nm (500–580 nm) as described previously ^58^. All confocal imaging was performed on a Leica TCS SP8 STED 3X confocal laser scanning microscope (Leica Microsystems) using a 63×/1.40 oil objective. Images of mitochondria (colocalization and morphology) were deconvoluted using Huygens Professional software.

### Seahorse metabolic assays on cells

Oxygen consumption rates (OCR) and extracellular acidification rates (ECAR) were measured using Seahorse XF96 bioscience extracellular flux analyzer (Agilent). Seahorse XF Cell culture microplates were treated with Cell Tak (Thermo Fisher Scientific) according to the manufacturer’s instructions and number of viable cells was determined using Countess Automated cell coulter with trypan blue stain before plated in 22 or more replicates at a density of 25000 cells/ well. For measurement of basal and maximal OCR myeloma KO and WT cells were incubated in XF assay medium supplemented with 10 mM Na-Pyruvate, 2 mM glutamine (both from Sigma) and 10 mM glucose (Merck) followed by injections of oligomycin, carbonyl cyanide p-trifluoro-methoxyphenyl hydrazone (FCCP); and antimycin A+ rotenone at final concentrations of 1 µM, 1 µM, and 2 µM +2 µM, respectively. For evaluation of glycolysis, cells were incubated in XF assay medium with only 2 mM glutamine, followed by injections of glucose, oligomycin and 2-deoxy-glucose (2-DG) at final concentrations of 10 mM, 1 µM and 50 mM, respectively. For IL-32 rescue cells, dead cells were removed by Optiprep^TM^ (Stemcell Technologies, Vancouver, Canada) according to manufacturer’s instructions and viable cells counted with the Countess Automated Cell Coulter before basal OCR and ECAR were evaluated by combining the mito stress and glycolysis stress test, also described elsewhere^10^. The same medium supplements as for mito stress assay were used, except glucose, which was added as the first injection in the assay (final concentration of 10 mM) before injections with oligomycin, FCCP and Rotenone+antimycin+ 2-DG. Basal and maximum OCR and basal ECAR were calculated according to manufacturer’s instructions. The OCR/ECAR ratio was calculated from basal OCR and basal ECAR values measured in the mito stress assays.

### Seahorse metabolic assay on isolated mitochondria

Mitochondria from 70-100*10^6^ IL-32 KO and WT cells were isolated as previously described^59^. Isolated mitochondria were quantified by Bradford assay and resuspended to desired concentration in mitochondrial assay buffer (MAS) (70mM sucrose, 220 mM mannitol, 10 mM KH_2_PO_4_, 5 mM MgCL_2_, 2 mM HEPES, 1 mM EGTA and 0.2% fatty acid free BSA) which had been supplemented with 2 mM malate, 10 mM Na-pyruvate and 5 mM glutamic acid and preadjusted to pH 7.2 at 37°C. 20 μg mitochondria were plated in each well of an XFe96 seahorse plate and the plate was spun at 2000 x g for 20 min at 4°C for attachment of the mitochondria. After centrifugation prevarmed (37°C) MAS + substrates were added to each well to a final volume of 180 μl incubated in a non-CO_2_ incubator for 20 min, before basal oxygen consumption rate (OCR) was measured using Seahorse XF96 bioscience extracellular flux analyzer (Agilent, CA, US).

### ROS and ATP quantification in cells

Mitochondrial ROS (mtROS) was assayed by MitoSoX Red staining, mitochondrial membrane potential and mitochondrial mass were quantified by TMRM and mitotracker green staining, respectively (all from Thermo Fisher). The procedures are described in more detail in Supplemental Methods. For ATP measurements, 40 000 cells were seeded/well and ATP levels were measured by CellTiter-Glo Luminescent (CTG) Cell Viability Assay (Promega) following manufacturer’s instructions. Luminescence was recorded using a Victor3 plate reader and Wallac 1420 Work Station software (PerkinElmer Inc.).

### Mitochondrial ROS and membrane potential in isolated mitochondria

Isolated mitochondria were quantified by Bradford assay and stained for 10 min at 37°C in a 20 % O_2_, 5% CO_2_ incubator with 5 μM MitoSox Red (Thermo Fisher) in MAS (70mM sucrose, 220 mM mannitol, 10 mM KH_2_PO_4_, 5 mM MgCL_2_, 2 mM HEPES, 1 mM EGTA and 0.2% fatty acid free BSA) which had been supplemented with 2 mM malate, 10 mM Na-pyruvate and 5 mM glutamic acid and preadjusted to pH 7.2 at 37°C. After staining the mitochondria were washed once in MAS and resuspended in MAS. 10 *u*g mitochondria/well were plated in 96-well optical plates followed by stimulation with 50 ng/mL rhIL-32*y* or rhIL-32β for 60 min 37°C (20% O_2_, 5% CO_2_ incubator). The fluorescence was assessed at with excitation 531 nm and emission 590 nm using Victor3 plate reader and Wallac 1420 Work Station software. For assessment of membrane potential, the isolated mitochondria were prepared in MAS buffer, stained with 500 nM Mitotracker Orange CMTMRos for 30 min 37°C in a 20% O_2_, 5% CO_2_ incubator before washed with MAS and seeded in 10 μg/well in 96-well optical plates and assessed immediately with excitation 531nm and emission 590 nm using Victor3 plate reader and Wallac 1420 Work Station software.

### RNA-sequencing of IL-32 KO cell lines

INA-6, JJN3 and H929 KO and WT cells were harvested from basal culture conditions and RNA was isolated using RNeasy kit (Qiagen). H929 and JJN3 cells were sequenced using the library preparation kit SENSE mRNA-Seq Library Prep Kit V2 from Lexogen, followed by 75bp single read sequencing on the NextSeq 500 Illumina machine. The sequencing depth was 5 million reads per sample. The INA-6 cells were sequenced (18 million reads per sample) using the TruSeq Stranded mRNA library preparation kit from Illumina (#20020595) followed by 75bp single read sequencing on the Illumina Hiseq 4000 next machine. IL-32 isoform analysis was performed using Kalisto (v0.43.0) using the parameter -b 100 and otherwise default parameters.

### RNA-sequencing data analyses

RNA sequencing data (MMRF_CoMMpass_IA13a_E74GTF_Salmon_Gene_Counts) and clinical data were downloaded from the Multiple Myeloma Research Foundation CoMMpass IA13 release. RNA sequencing data from CD138^+^cells were available for 795 baseline samples from patients with MM. Data on overall survival and progression-free survival were available for all these patients. RNA-sequenced CD138+ cells from longitudinal samples were available for 47 samples in IA13. We analyzed IL-32 expression in 47 patients at diagnosis and first relapse timepoint. For survival analyses, patient samples taken at diagnosis were divided into high and low IL-32 expression based on the upper 10th percentile (n = 54; counts per million (cpm,log2)> 1.52) and lower 90th percentile (n = 741, (cpm,log2)< 1.52). Differentially expressed genes between high (10th percentile) and low IL32 (90th percentile) were assessed using the same percentiles, and expression-requirement of 1cpm in at least 20% of the samples. GO analyses were performed using R package ClusterProfiler (v3.14.3) using expressed genes as background. Cutoff for significance of genes implemented in GO was p=0.01. The GO-analyses for the IL32-KO and WT sequencing data were performed as described above for the CoMMpass data (see figure texts for filtering of data). Survival-analysis was performed in R, using the package “survival”. High and low IL-32 was defined as previously described and “Time” and “Status/Sensoring” were collected from the clinical data in ComMMpass. Survival curves were plotted using “ggsurvplot” in R using the package “survminer”.

Differentially expressed genes were calculated using limma-voom in R. We required genes to be expressed with at least 1 cpm in 20% of the samples. We used TMM normalization for calculating the “calcNormFactors” in limma. P-values were adjusted using Benjamini-Hochberg.

All analyses were run using R version 3.6.2 (2019-12-12). Used packages with version number includes:packageVersion(“survminer”)‘0.4.6’;packageVersion(“biomaRt”)‘2.41.4’;packageVersion(“clusterProfiler”)‘3.12.0’;packageVersion(“org.Hs.eg.db”)‘3.8.2’;packageVersion(“limma”)‘3.40.6’;packageVersion(“edgeR”)‘3.26.8’;packageVersion(“survival”)‘3.1.8’; packageVersion(“pheatmap”)‘1.0.12’; packageVersion(“ggplot2”) ‘3.2.1’.

### Single-cell transcriptome analysis

The single cell data was download from Gene Expression Omnibus (GEO) using the accession number GSE106218. The data was analyzed as described by Ryu et al. ^28^ using the *Seurat* package in *R.* ^60^ Specifically, dimension reduction was performed using uniform manifold approximation and projection (UMAP) by using the 10 first principal components from the *FindNeighbors* and *FindClusters* functions in *Seurat*. IL-32-expressing cells were defined as cells for which at least one read for IL-32 was detected and IL-32 non-expressing cells were defined as cells for which no reads for IL32 were detected. High and low IL32 expression is defined similarly. Differentially expressed genes between IL-32-expressing cells and IL-32 non-expressing cells were detected using *limma-voom* ^61^ as previously described for single cell data.^62^ Gene ontology (GO) analysis was performed using the package *clusterProfiler Yu* ^63^ and the *enrichGO* function in *R*. The p-values were adjusted using Benjamini-Hochberg method and a q-value cutoff of 0.05 was used. Genes with log2 fold change above 1.5 was used as input in the gene ontology analysis and genes expressed in at least one cells were used as background. The *simplify* function in *clusterProfiler* was used to merge similar GO-terms. The GO-terms were order by q-value and the top 20 terms were plotted using *ggplot* in R.

### Flow cytometry

For assessment of CD38 and CD45 on IL-32 KO and WT cells, live cells were stained with anti-CD38 PE-Cy7 (#335825), CD45-FITC (#555482) antibodies from BD biosciences and assessed by flow cytometry. Samples were analyzed with FlowJo V10.

### Quantification of immunoglobulin kappa light chain

1*10^6^ cells were seeded in 1 ml RPMI, 0.1% BSA with the addition of 1 ng/ml IL-6 for INA-6 cells and incubated for 24 hours in 5% (normoxia) or 2% (hypoxia) O_2_. The cells were counted, and Ig kappa light chain concentration were quantified in the cell culture supernatant by ELISA according to the manunfacturer’s protocol (Human Kappa ELISA, Bethyl Laboratories).

### Statistics

Statistical analyses were performed using GraphPad Prism version 9 (GraphPad Software) unless otherwise stated. Paired or paired ratio or unpaired Student’s t-test or Wilcoxon signed-rank test were used to compare two groups. For comparison of two groups, and more than two groups with measurements over time, multiple T-tests and two-way ANOVA followed by Sidàk’s or Dunnett’s multiple comparisons test was used, respectively.

## Data sharing statement

For original data, please contact.

## Acknowledgements

The authors thank Berit Størdal and Hanne Hella for technical support and Pegah Abdollahi for help and training in experimental procedures. The confocal imaging services were provided by the Cellular and Molecular Imaging Core Facility (CMIC), Norwegian University of Science and Technology (NTNU), which is funded by the Faculty of Medicine at NTNU and Central Norway Regional Health Authority. We thank previous and current members of our patient advisory board and the Multiple Myeloma Research Foundation Personalized Medicine Initiatives (https://research.themmrf.org and www.themmrf.org) which generated the CoMMpass data.

This work was supported by funds from the Norwegian Cancer Society (#206643, #191861), the Research Council of Norway (#223255), the Liaison Committee for education, research and innovation in Central Norway (# 90061000 and #30171600) and the Cancer Fund at St. Olavs Hospital, Trondheim, Norway.

## Author contributions

K.R.A. designed and performed experiments, analyzed and interpreted data and wrote the paper, R.M., M.H.K., S.S.T. and L.M.B. performed experiments, analyzed and interpreted data, M.W., M.Z., S.H.M. G.B, A.M.S., A.S and RG contributed to experiments and data collection, R.M and K.M. performed bioinformatics analyses, T.S.S. and A.W. contributed with patient clinical data and T.S. initiated and designed the study, analyzed and interpreted data and wrote the paper. All authors revised the manuscript.

