## Supplementary Figures for "IL-32 is a metabolic regulator promoting survival and proliferation of malignant plasma cells"

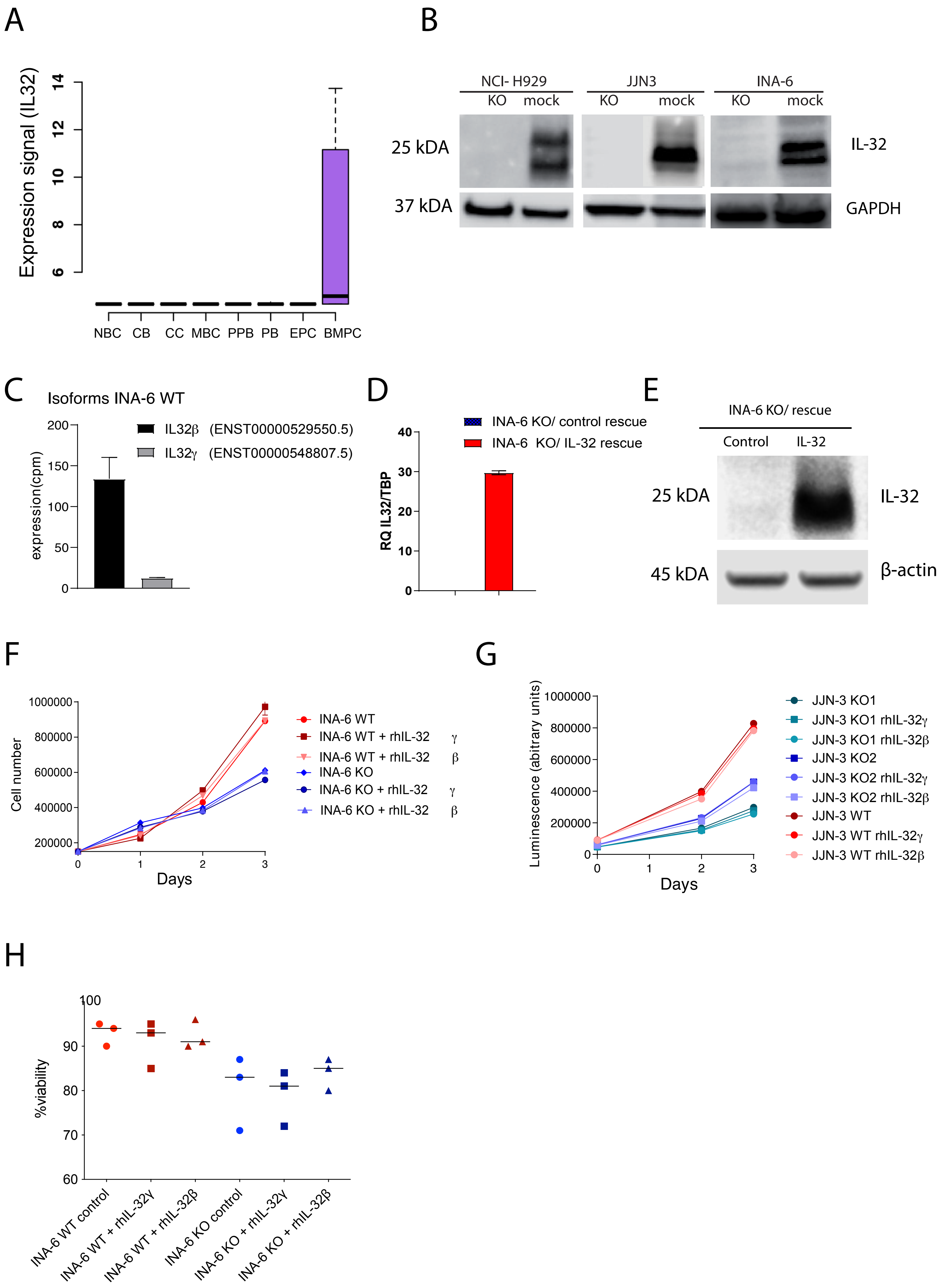


Supplementary Figure 1

**Supplementary Figure 1. IL-32 expression.**

**(A)** IL-32 gene expression analyzed in Affymetrics microarray data (GCRMA normalized) of B cell subtypes derived from healthy individuals publicly available at GenomicScape.com (ID: GS-DT-2). NBC: naïve B cells (n=5), CB: centroblasts (n=4), CC: centrocytes (n=4), MBC: memory B cells (n=5), PPB: preplasmablasts (n=5), PB: plasmablasts (n=5), EPC: early plasma cells (n=5), BMPC: bone marrow plasma cells (n=5).

**(B)** H929, JJN-3 and INA-6 IL-32 KO cells were generated by CRISPR/Cas9 gene editing, using lentiviral transduction and plasmid transfection, respectively. Efficiency of *IL32* knockout was evaluated by western blotting of IL-32.

**(C)** IL-32 isoforms in RNA-sequenced INA-6 WT mock cells. The Ensembl transcript ID is shown in brackets.

**(D)** INA-6 KO IL-32 rescue cells were generated by transducing IL-32 KO cells with an IL-32 lentiviral vector and knock-in efficiency evaluated by western blot of IL-32.

**(E)** IL-32 mRNA in INA-6 KO IL-32 rescue cells assessed by qPCR. Mean± SD of technical replicates of one representative of 3 experiments is shown.

**(F)** INA-6 KO and WT mock cells were treated with rhIL-32 isoforms α, β and γ and proliferation was evaluated by cell counting for 3 days. Each time point shows the mean ± SD of two technical replicates.

**(G)** JJN-3 KO and WT mock cell proliferation with and without medium supplementation of rhIL-32 isoforms was assessed by CTG assay. Arbitrary units were normalized to values at day 0 for each cell line. Each time point shows the mean ± SD of 3 technical replicates.

**(H)** Viability of INA-6 KO and WT mock cells treated with rhIL-32 isoforms α, β and γ overnight evaluated by trypan blue staining. The mean ± SEM for 3 independent experiments is shown. Statistical significance was evaluated by two-way ANOVA and Dunnett´s multiple comparisons test for F-H. There were no significant differences.

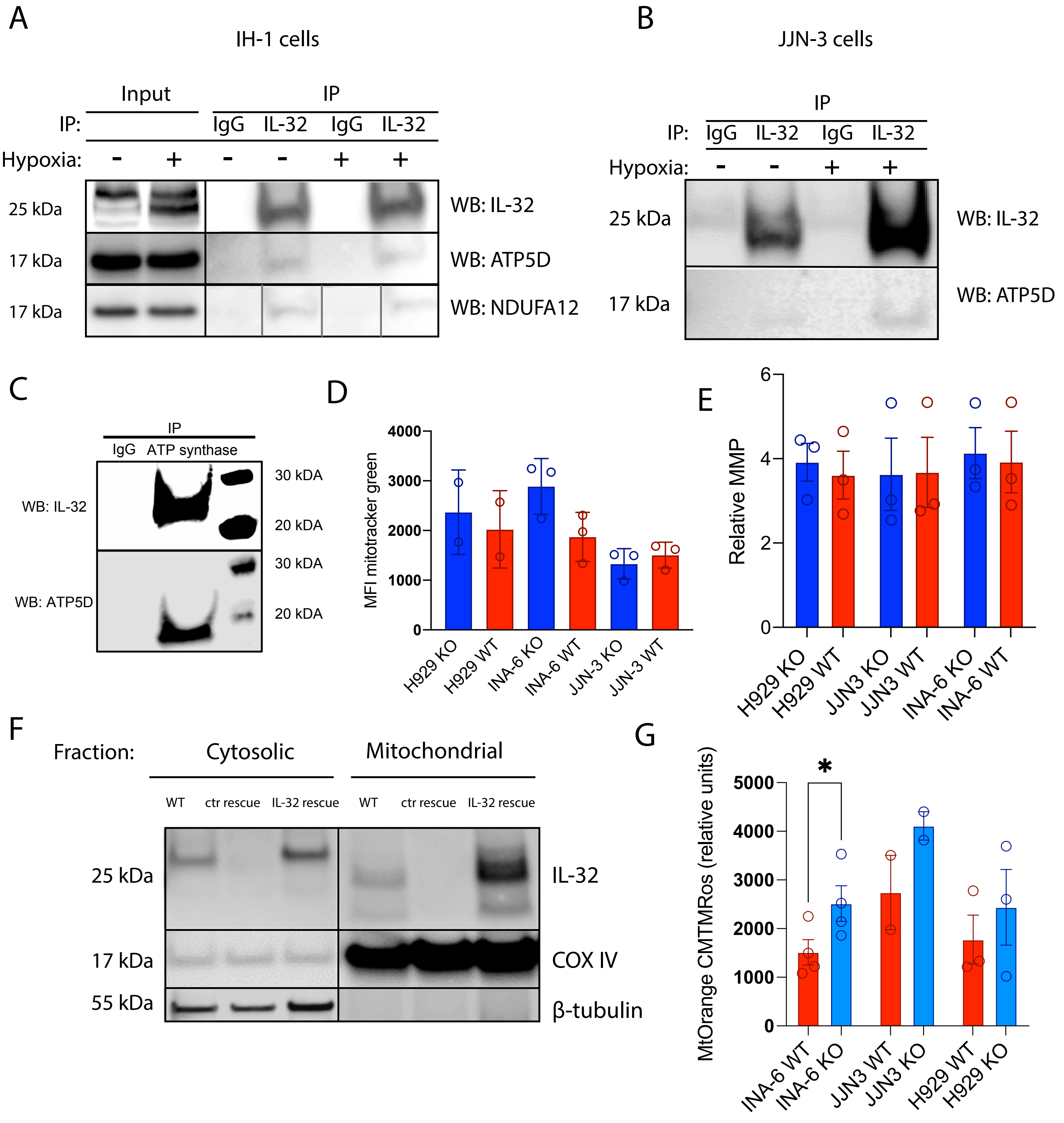


Supplementary Figure 2

**Supplementary Figure 2:**

**(A)** CO-IP was performed by pulldown of endogenous IL-32 from IH-1 cells, and ATP5D, NDUFA12 and IL-32 was detected by western blotting. For NDUFA12 the order of "IP: IgG hypoxia" and "IP: IL-32 no-hypoxia" was switched as indicated by grey lines.

**(B)** CO-IP was performed by pulldown of endogenous IL-32 from JJN-3 cells, and ATP5D and IL-32 was detected on western blot. The western blot shown is one of two independent experiments.

**(C)** CO-IP was performed by pulldown of the ATP synthase complex in INA-6 cells and IL-32 and ATP5D was detected by western blot. Figure shows one representative WB of 3 independent experiments.

**(D)** Mitochondrial mass in IL-32 KO and WT cell lines quantified by mitotracker green**.** Figure shows the mean ± SEM of 3 independent experiments. The differences were not significant.

**(E)** Relative mitochondrial membrane potential (MMP) in myeloma IL-32 KO and WT cell lines quantified by TMRM/ mitotracker green flow cytometry. Figure shows the mean ± SEM for 3 independent experiments. For INA-6 only one independent experiment was performed for KO2. There were no significant differences.

**(F)** Western blot of IL-32 in mitochondrial and cytosolic fraction of INA-6 IL-32 rescue cells and rescue control cells.

**(G)** Mitochondrial ROS in isolated mitochondria myeloma KO and WT cell lines quantified by MitoSox Red. Figure shows the mean ± SEM of biological replicates.

P-values in D, E, G and H were analyzed by unpaired Student´s t-test. * $p \leq 0.05$, *** $p \leq 0.001$, **** $p \leq 0.0001$.

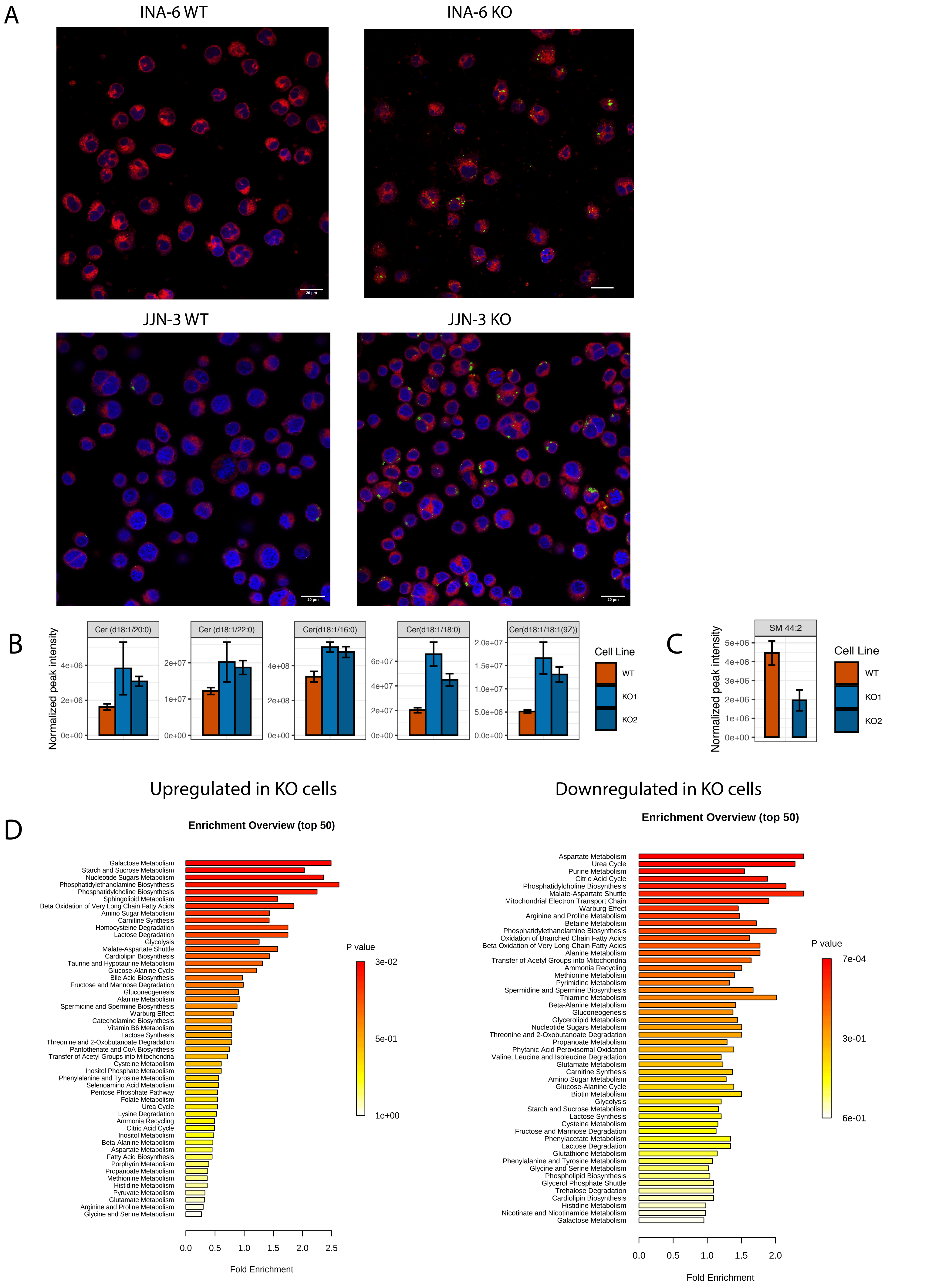


Supplementary Figure 3

**Supplementary Figure 3:**

**(A)** Lipid droplets in INA-6 and JJN-3 KO and WT mock cells, stained with Nile Red. Polar lipids (red) were excited at 590 nm (600–700 nm) and neutral lipids (green) at 488 nm (500–580 nm). Confocal imaging was performed with a Leica TCS SP8 STED 3X using a 63X 1.4 (oil) objective and LAS X software. Scale bar: 20 μM.

(**B)** Significantly (P<0.05) altered ceramide species between INA-6 KO1, KO2 and WT mock cells.

**(C)** Significantly (P=0.00042571) altered sphingomyelin (44:2) betweenINA-6 KO2 and WT mock cells. Metabolite absent/ not detected in KO1.

**(D)** Pathway enrichment of significantly downregulated and upregulated metabolites in INA-6 KO cells compared to WT cells using Metaboanalyst.

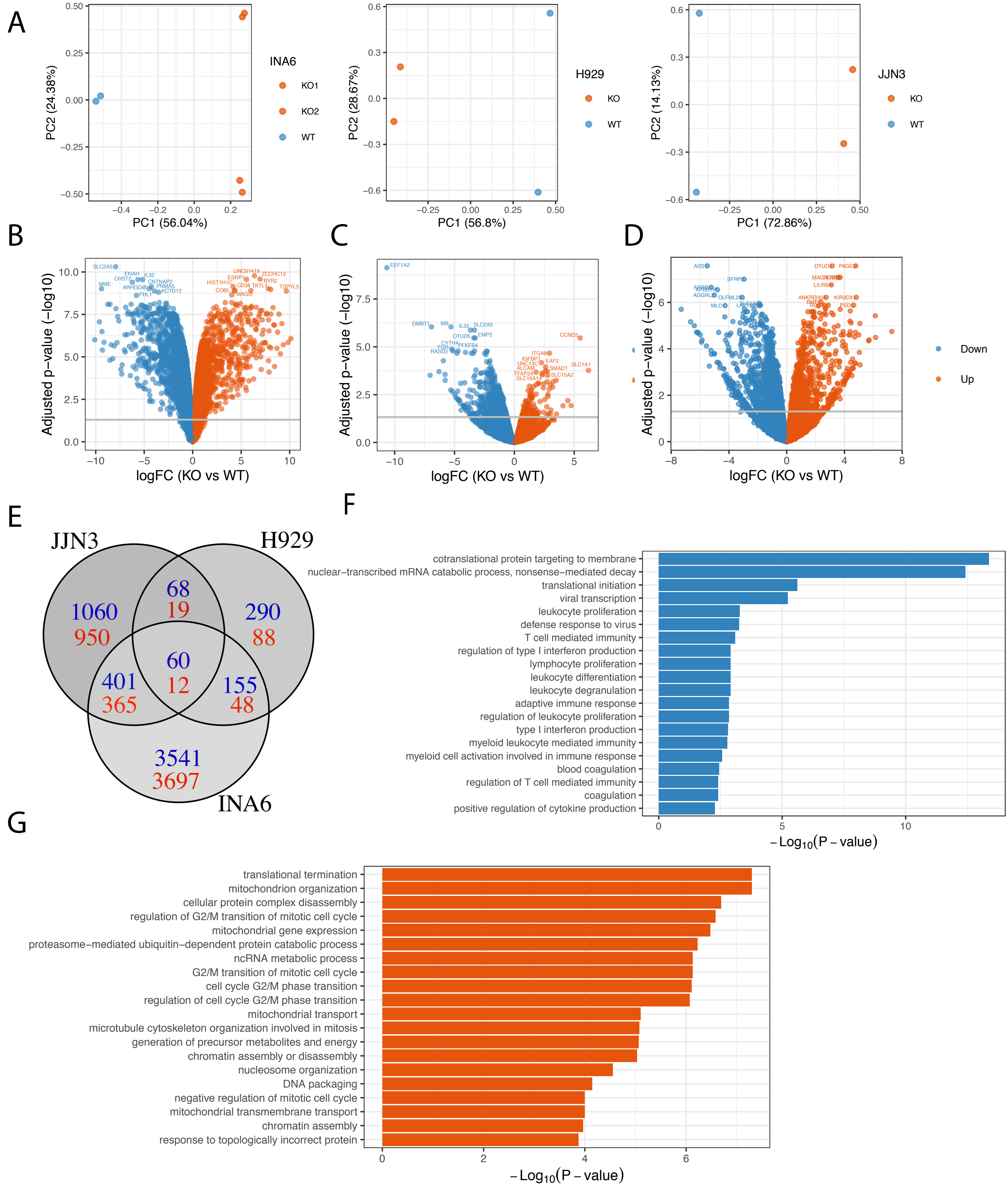


Supplementary Figure 4

**Supplementary Figure 4: RNA-sequencing of IL-32 KO cell lines**

**(A)** Two INA-6 KO cell lines and WT mock cells, a H929 and JJN-3 KO cell line and WT mock cells were subjected to RNA sequencing and the PCA plots show the overall differences in gene expression between KOs and WT mock.

**(B-D)** Volcano plot showing the most significantly upregulated and downregulated genes in (B) INA-6 KO cells (2 clones) (C) H929 KO cells and (D) JJN-3 KO cells vs WT mock cells. Statistical significance analyzed by limma in R with Benjamini-Hochberg-adjusted P-values.

**(E)** Venn-diagram of the shared upregulated (red) and downregulated (blue) genes in IL-32 KO myeloma cell lines, using a cutoff of FC>1 for upregulated genes and FC<-1 for downregulated genes.

**(F, G)** GO-analysis of the differentially expressed genes (Benjamini-Hochberg-adjusted P-value < 0.05; log2 fold change >0 and <0 for up- and -down-regulated genes, respectively) between INA-6 KO and WT mock cell lines. The figures show the top 20 significantly enriched biological processes downregulated **(F)** or upregulated **(G)** in KOs vs WT mock. The GO terms are ordered by the Benjamini-hochberg adjusted P-values.

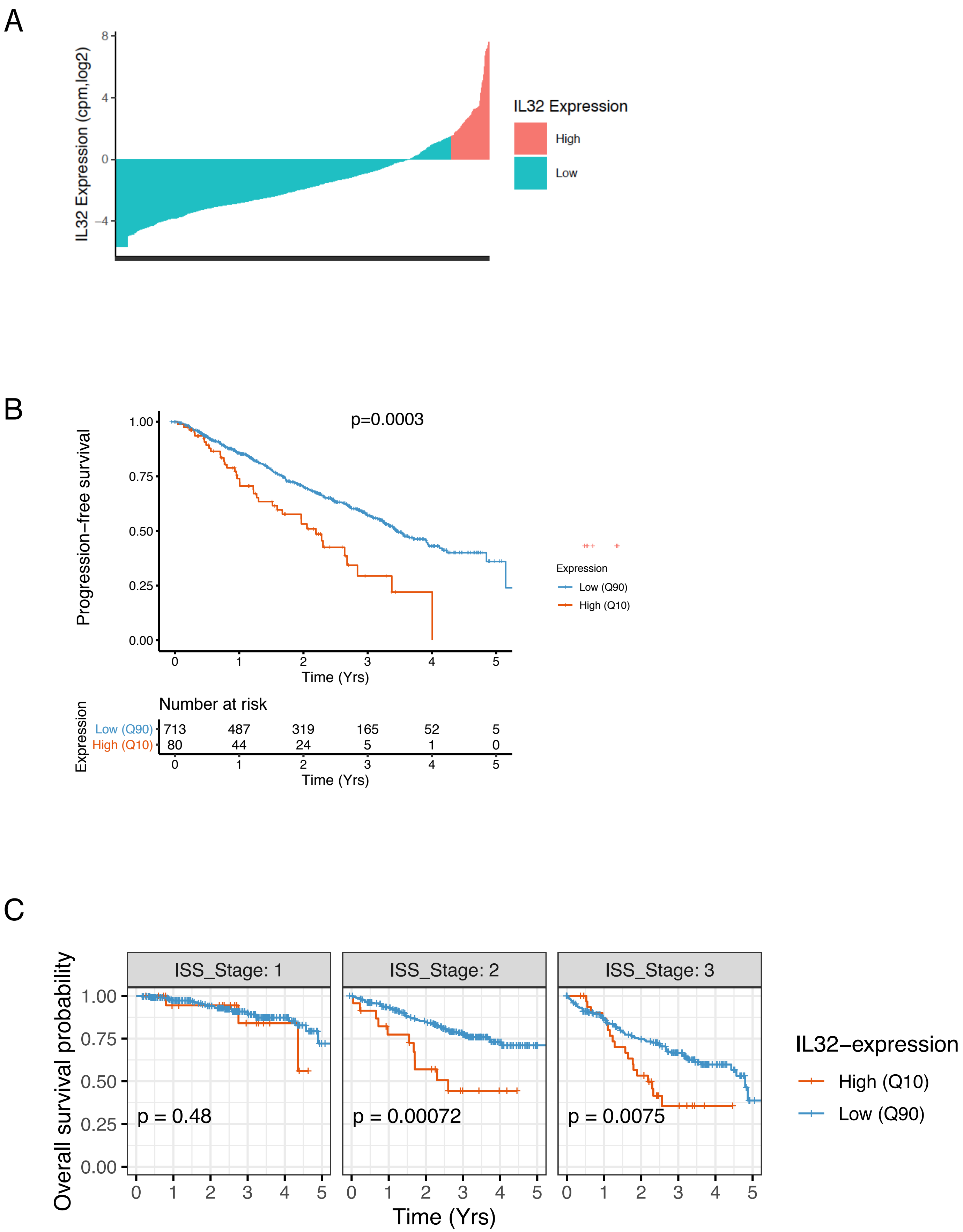


Supplementary Figure 5

**Supplementary Figure 5. IL-32 is a prognostic factor**

**(A)** IL-32 expression in all patients from CoMMpass IA13 (N= 792), upper 10th percentile (n= 80, log2 cpm> 1.52) colored in pink**.**

**(B)** Progression- free survival of IL-32 expressing patients ($10^{th}$ percentile) compared to non-expressing patients ($90^{th}$ percentile) in the IA13 CoMMpass dataset p= 0.0003, using Cox proportional-hazards regression model.

**(C)** Assessment of IL-32 as an independent prognostic factor for overall survival in IA13. IL-32 expressing patients ($10^{th}$ percentile) compared to non-expressing patients ($90^{th}$ percentile) at different ISS stages (stage I, stage II, or stage III disease). Adjustment for ISS stage was performed using multivariate Cox-regression, using the coxph function in R.

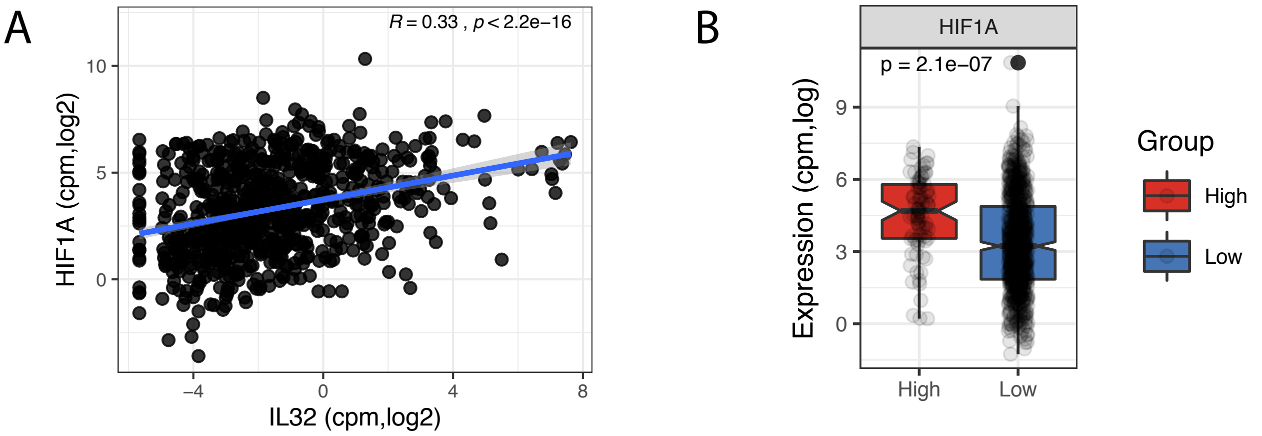


Supplementary Figure 6

**Supplementary Figure 6. IL-32 expression correlates with *HIF1A***

**(A)** Correlation between IL-32 and *HIF1A* in CoMMpass IA13, evaluated by Pearson correlation coefficient.

**(B)** *HIF1A* gene expression in IL-32-expressing patients (upper $10^{th}$ percentile) compared to non-expressing patients (lower $90^{th}$ percentile) in CoMMpass IA13. Significance was determined by two-tailed Wilcoxon signed-rank test.

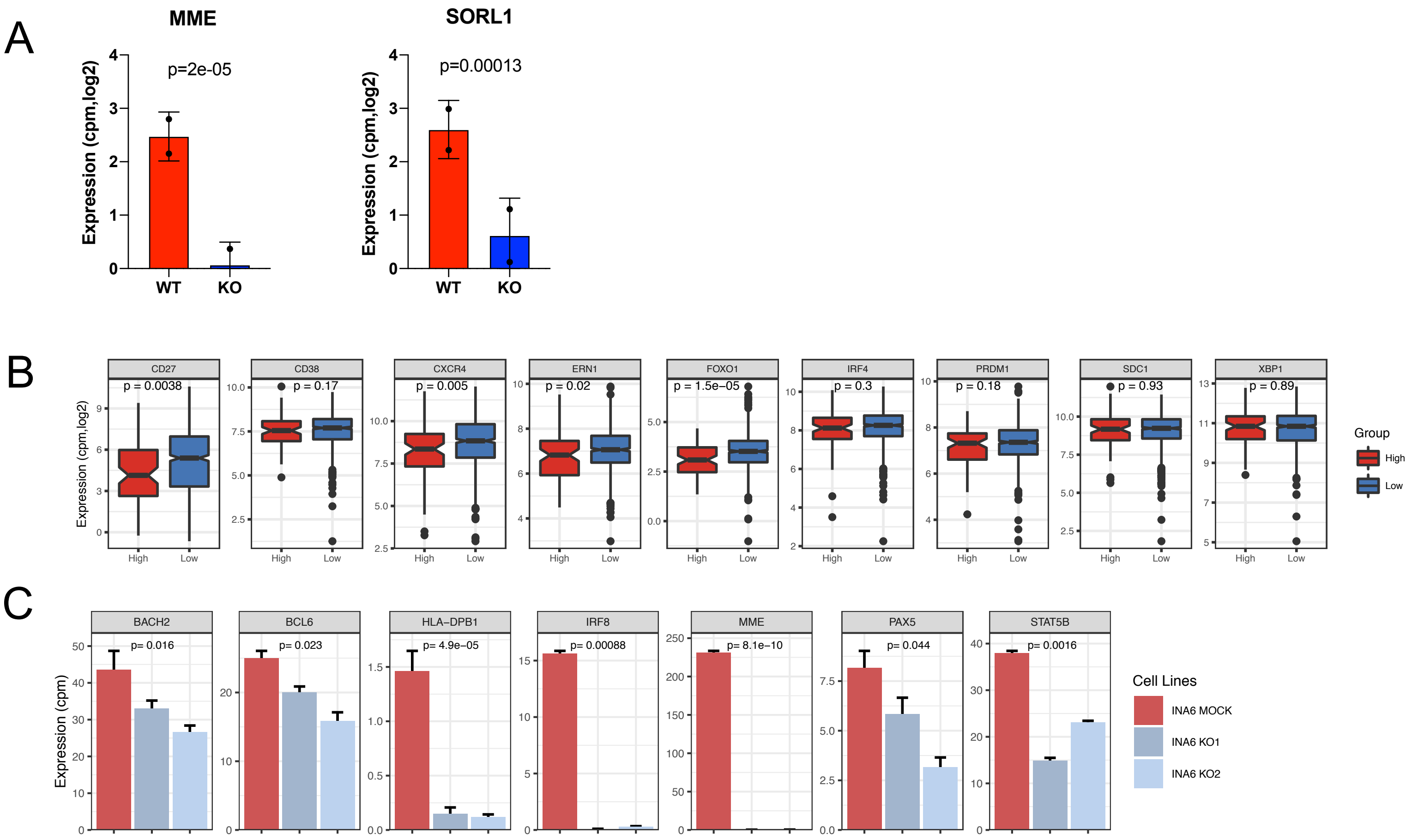


Supplementary Figure 7

**Supplementary Figure 7. IL-32 expression is associated with an immature phenotype and carfilzomib resistance.**

**(A)** Gene expression of *MME* and *SORL1* in H929 IL-32 KO and WT mock cells. significance calculated by limma t-test with Benjamini-Hochberg-adjusted p-values.

(**B)** Evaluation of gene expression of typical mature plasma cell markers (based on a literature search) in IL-32 expressing patients (upper10$^{th}$ percentile) compared to non-expressing patients (lower 90$^{th}$ percentile) in CoMMpass IA13. Significance analyzed by two-tailed Wilcoxon signed-rank test.

(**C)** Genes associated with less differentiated stages of B-cell maturation downregulated in INA-6 KO cells assessed from RNA-sequencing data of INA-6 KO1, KO2 and WT mock cells. P-values analyzed by limma T-test.
